# Telmisartan alters the transcriptomes of cancer associated fibroblasts and fibrocytes in a solid tumour model

**DOI:** 10.64898/2026.09.12.751091

**Authors:** Che-Min Lee, Nikita Telkar, Brennan J Wadsworth, Rachel A Cederberg, Rocky Shi, Lisa Zhan, Prabhreet Sekhon, Meredith Clark, Michael GS Hall, Kiersten N Thomas, Wan L Lam, Kevin L Bennewith

**Author notes:** **Corresponding Author:** Dr. Kevin L Bennewith, PhD Distinguished Scientist, BC Cancer Research Institute #10-108, 675 West 10^th^ Ave, Vancouver, British Columbia, Canada, V5Z 1L3.

## Abstract

Cancer associated fibroblasts (CAFs) are the main contributors to fibrosis in solid tumours. Fibrosis can lead to increased tumoural stiffness, immune cell exclusion, and poor response to cancer therapies. However, therapeutically depleting CAFs has been shown to increase tumour progression and metastasis in some pre-clinical tumour models. Targeted approaches to inhibit CAF activity and decrease fibrosis without abrogating CAFs altogether may improve the effect of existing cancer therapies. We have previously shown that the blood pressure medication telmisartan decreases tumoural collagen, resulting in improved vascular perfusion, decreased hypoxia, and improved radiation treatment efficacy. To understand how telmisartan influences CAF populations in solid tumours, we conducted single cell RNA sequencing analyses of host cells isolated from tumours in mice treated with telmisartan. Telmisartan modified the transcriptomes of CAF populations, promoting the expression of chemokine signaling while decreasing the expression of extracellular matrix components in myCAFs. Telmisartan decreased expression of collagen transcripts in myCAFs and fibrocytes, indicating that telmisartan does not remove myCAFs but changes the myCAF transcriptome to become less fibrillar. Our work provides further support for using telmisartan to manipulate CAF populations in the solid tumour microenvironment to decrease collagen deposition and render tumours more responsive to current cancer therapies.

## Introduction

Fibrosis or desmoplasia in solid tumours is caused by the deposition and remodeling of dense collagen matrices that are associated with diminished responses to many types of cancer therapies^1–3^, and ultimately poor patient outcomes^4,5^. Aberrant collagen production from cancer-associated fibroblasts (CAFs) is well-established, although the recent identification of monocyte-derived fibrocytes in solid tumours may also contribute to tumoural fibrosis and dense extracellular matrix (ECM)^6,7^. CAF subsets can include myofibroblast-like myCAFs, inflammatory iCAFs, and antigen-presenting apCAFs. The myCAFs are defined by expression of the myofibroblast contractile protein alpha smooth muscle actin (αSMA; transcript *Acta2*), high levels of ECM secretion and remodelling components, and contribute the most to ECM density in solid tumours^8,9^. Conversely, iCAFs are mainly defined as cells that secrete cytokines and recruit immune cells (particularly myeloid cells) to the tumour. Fibrocytes are a poorly studied cell type that have previously evaded identification in tumour studies. Fibrocytes are known to be monocyte-derived, but have both ECM secretion and antigen-presentation capabilities. Fibrocytes are difficult to study as lineage tracing studies have shown that fibrocytes can lose expression of CD45 while retaining expression of myeloid cell markers^7,10^. Additionally, fibrocytes are also defined by the expression of fibroblast-related genes such as collagens and other ECM components. Preclinical tumour models that contain myCAFs, iCAFs, and fibrocytes better allow us to study how these cell types contribute to ECM deposition and ultimately how tumours respond to cancer therapies^11–13^.

MyCAFs are predominantly responsible for producing aberrant levels of collagen in solid tumours, and one class of drugs that shows promise in targeting myCAF-related fibrosis is the angiotensin II type 1 receptor (AT1R) blockers (ARBs). ARBs are commonly prescribed anti-hypertensive medications that have also been shown to exhibit anti-fibrotic properties. Angiotensin II (Ang II) signaling in fibroblasts is known to cause collagen deposition and fibrosis, thus ARBs that block Ang II binding to AT1R can act as anti-fibrotics in different pathological conditions including solid tumours. The ARB losartan has been shown to decrease TGFβ1 signaling and collagen I (Col1) deposition in murine pancreatic ductal adenocarcinoma (PDAC) models, decompressing tumour blood vessels, improving vascular perfusion, and subsequently potentiating improved chemotherapy drug delivery^14^. Losartan is currently being tested in a phase II clinical trial as a neoadjuvant with FOLFIRINOX chemotherapy in patients with locally advanced PDAC^15^. Telmisartan is a newer generation ARB with longer bioavailability and increased lipophilicity and AT1R binding affinity than losartan^16^. We previously showed that telmisartan can significantly decrease collagen deposition in human tumour xenografts, leading to increased vascular perfusion and decreased hypoxia that improved response to ionizing radiation therapy^17^. However, the influence of telmisartan on different CAF populations and fibrocytes in tumours is unknown and may have important implications for applying telmisartan to modulate the solid tumour microenvironment to improve cancer therapy. Herein, we use flow cytometry and single cell RNA sequencing to identify how telmisartan influences cellular phenotypes in a human colorectal adenocarcinoma tumour xenograft model with a particular focus on CAF subsets.

## Results

To understand how telmisartan influences CAFs in WiDr xenograft tumours, we first evaluated CAF response to telmisartan-treated WiDr tumours via flow cytometry. Telmisartan was provided *ad libitum* in the drinking water of mice implanted with WiDr tumours, and the tumours were harvested 44 days after implant at a size of 400-500 mg. After processing tumours into a single cell suspension and gating for live singlets, CAFs were identified as CD45^-^EpCAM^-^CD31^-^ (**Figure 1A**, **Supplementary Figure 1A**). Within those cells, double-positive PDPN^+^CD90^+^ cells were identified as active CAFs (**Figure 1B**). Two CAF populations were described based on Ly6C and CD29 surface expression within active PDPN^+^CD90^+^ CAFs as follows: inflammatory iCAFs as Ly6C^hi^ CD29 ^mid^ and myofibroblast myCAFs as Ly6C^mid-low^ CD29^mid-hi^ (**Figure 1B, Supplementary Figure 1B**). Myofibroblasts are often categorized by αSMA expression, and we found that αSMA and CD29 expression were highly similar (**Figure 1B**). Antigen presenting CAFs (apCAF)s are often defined as CD74^+^ MHC II^+^, although we found no detectable MHC II (IA/IE) expression in CAFs (**Supplementary Figure 1C**) and observed only 1-2% of active CAFs expressing CD74 (**Supplementary Figure 1D**). Thus, apCAFs were not explored further in this study. Within the active CAFs, telmisartan treatment decreased the myCAF population compared to control tumours, but did not affect the iCAF population frequency (**Figure 1C**). Additionally, telmisartan treatment resulted in decreased αSMA expression in the myCAFs based on decreased αSMA mean fluorescence intensity (MFI), while αSMA was not affected in the iCAFs (**Figure 1D**). Though telmisartan decreased the myCAF population in WiDr tumours, telmisartan did not affect the proportion of active PDPN^+^CD90^+^ CAFs, the proportion of live cells isolated from the tumours, or the tumour weight or mouse weight at endpoint (**Supplementary Figure 1E-H**).

**Figure 1.**
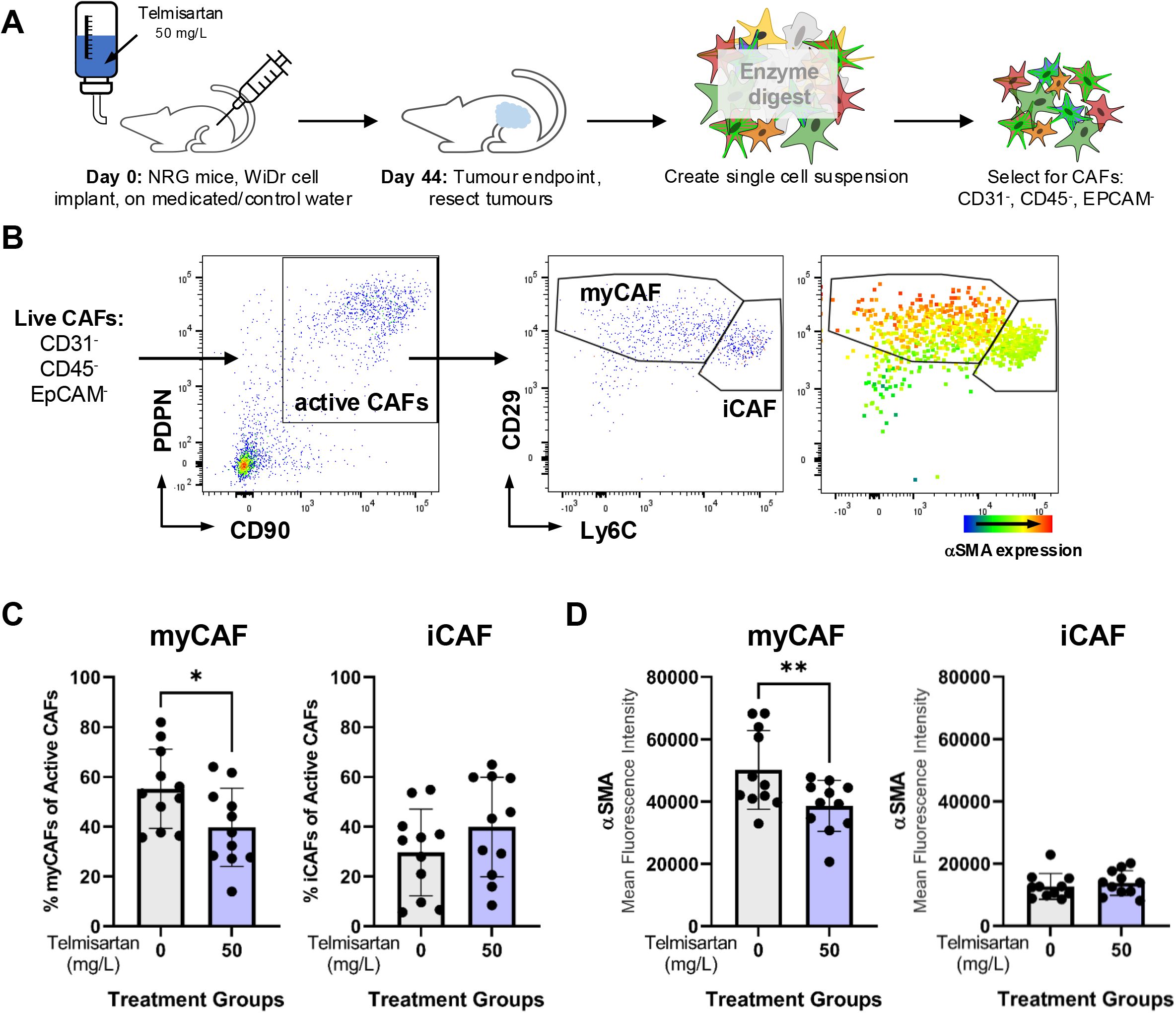
Heterogeneous CAF populations in WiDr tumours change with 50 mg/L telmisartan treatment. **A)** Mice were implanted with WiDr cells subcutaneously and put on telmisartan 50 mg/L water or acidified control water at the same time as implant. Tumours were taken when they reached around 500 mm^3^ (44 days for this experiment). Tumours were processed to a single cell suspension and CAFs were identified as live, singlet cells that were mouse CD31^-^, mouse CD45^-^, human EpCAM^-^. **B)** CD90^+^ PDPN^+^ CAFs distinguished as active CAFs which were further classified as myofibroblast myCAF (CD29^hi^, Ly6C^low-mind^) or iCAF (CD29^mid-hi^, Ly6C^hi^). Expression of αSMA overlaid on the CD29-Ly6C plot showed that high αSMA expression was present in myCAFs. **C)** myCAF and iCAF proportion of active CAFs in control or telmisartan-treated tumours. **D)** Mean fluorescence intensity (MFI) of myofibroblast marker, αSMA, in myCAF and iCAF cells.

We next utilized single-cell RNA-sequencing (scRNA-seq) to generate a higher resolution analysis of how telmisartan influences different cell types in WiDr tumours. Untreated control (CTRL) and telmisartan-treated (TEL) WiDr tumours were processed into single-cell suspensions and human EpCAM-expressing tumour cells were removed through magnetic bead separation to focus our scRNA-seq analysis on murine host cells from the tumours (**Figure 2A**). After quality control was conducted in the Seurat R package (**Supplementary Figure 2A-D**), the resultant 44,415 mouse cells were clustered via Louvain clustering into a UMAP projection at a resolution of 0.1, which defined 9 clusters (**Figure 2B**) and changes to the proportion of each cluster in CTRL vs TEL tumours (**Figure 2C)**. Clusters were identified using transcript expression of known cell markers (**Figure 2D-E, Supplementary Figure 2E**). Immune cells that were *Ptprc*+ (CD45 transcript) included neutrophils (cluster 1; *S100a8+, S100a9+, G0s2+*) and macrophages (cluster 2; *Ctss+, Fcgr1+, Ifi30+*). A fibrocyte population (cluster 3) was defined by myeloid cell markers (*S100a8, S100a9, Cd14, Cd74,* and *H2-Ab1*), low expression of *Ptprc* (CD45), and expression of fibroblast markers (*Col1a1, Acta2, Col3a1).* CAFs (*Col1a1+*) were segregated into myCAFs (cluster 4; *Acta2+* [αSMA], *Spp1+*, *Tnc+*) and iCAFs (cluster 5; *Ly6c+, Dcn+, Efemp+*). Cell populations present at lower proportions included keratinocytes (cluster 6; *Lgals7+*, *Sfn+*, *Perp+*), endothelial cells (cluster 7; *Pecam+, Plvap+, Egfl7+*), pericytes (cluster 8; *Rgs5+, Itga1+, Notch3+, Col1a1+*, *Acta2+*), and muscle cells (cluster 9; *Tnni2+, Acta1+, Des+*). We found that telmisartan treatment modified the proportions of normal cells in the tumours, decreasing the proportion of myCAFs and iCAFs with concomitant increases in the proportions of neutrophils and fibrocytes (**Figure 2C**). The ARB losartan can decrease thrombospondin (*Thbs1*) expression and TGFβ activity in murine tumours ^47,49^. Consistently, we found telmisartan-treated WiDr tumours contained myCAFs and fibrocytes with decreased expression of *Acta2*, *Thbs1*, the TGFβ target gene *Ccn2*, and the collagens *Col1a1* and *Col3a1* (**Supplementary Figure 2F**).

**Figure 2.**
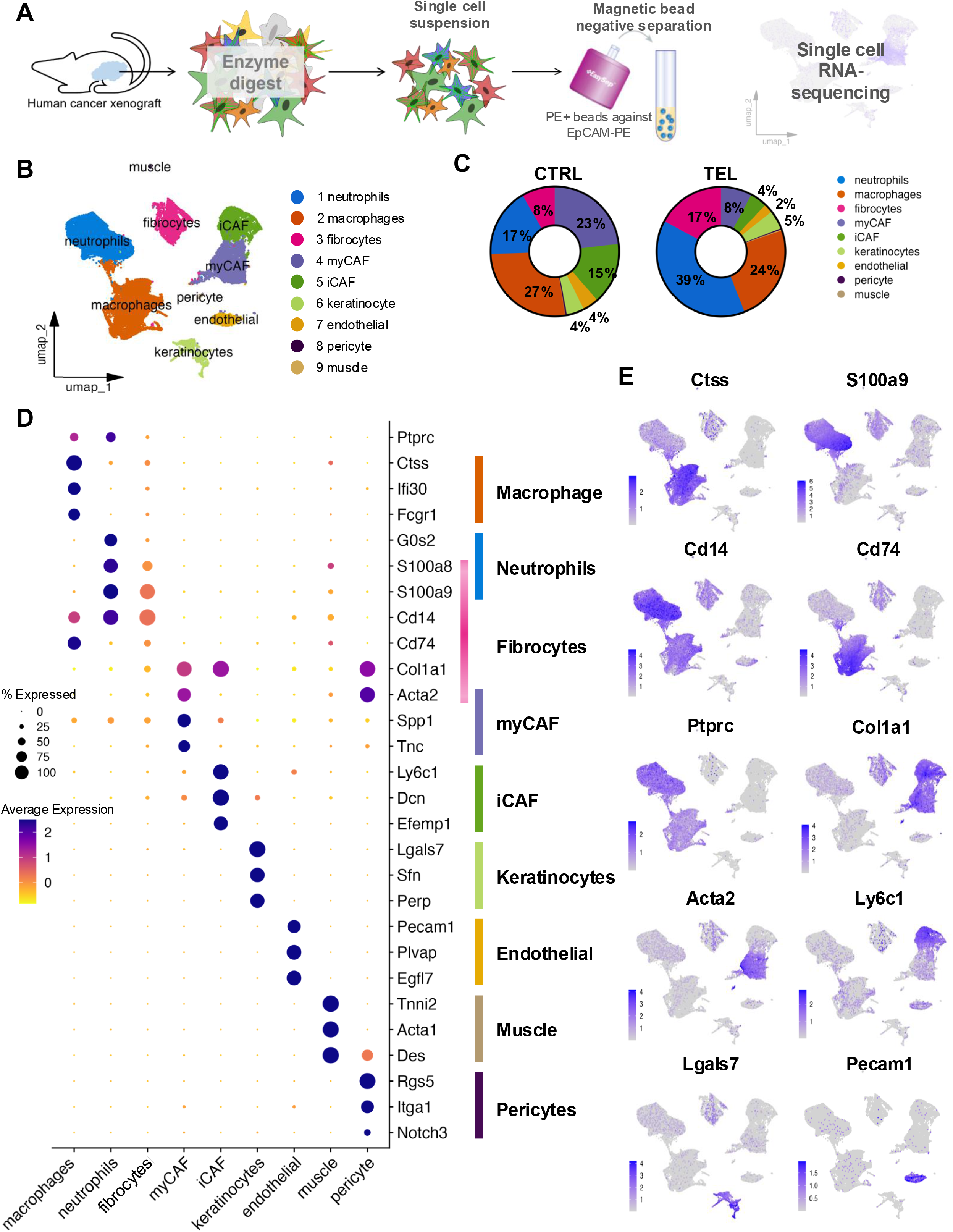
Single-cell RNA-sequencing of WiDr tumours with telmisartan treatment. **A)** Workflow to enrich for tumour microenvironment cells via magnetic bead separation in preparation for single-cell RNA-sequencing of WiDr tumours. **B)** UMAP projection of mouse cells from WiDr tumours clustered at resolution 0.1 with Louvain clustering to identify 9 clusters (0 -9) to be neutrophils (0), macrophages (1), myCAFs (2), fibrocytes (3), iCAFs (4), keratinocytes (5), endothelial cells (6), pericytes (7), and muscle cells (8). **C)** Single markers plotted on separate UMAPs that were used to define cell types. **D)** Dot plot of various markers used to define clusters.

Since fibrocytes have only recently been identified in solid tumours, we wanted to validate the scRNA-seq-based identification of fibrocytes by qPCR and flow cytometry. Circulating fibrocytes can be defined by myeloid cell markers such as CD45, CD11b, Ly6C, CD14, and S100A8/A9. When fibrocytes in tissues begin to produce extracellular matrix components, the cells express fibroblast markers including Col1, Col3, and αSMA, and can decrease expression of myeloid lineage markers including CD45 and CD11b. The fibrocyte population we identified through scRNA-seq had moderate expression of both myeloid and fibroblast cell markers (**Figure 2D-E**). CD14 expression in mice is distinct from CD14 in humans, where it is commonly used as a monocyte marker. As we saw moderate levels of *Cd14* expression in the fibrocyte population, we used surface CD14 protein expression to validate the WiDr tumour fibrocyte population by flow cytometry. Fibrocytes were identified as EpCAM^-^CD31^-^CD11b^-^CD90^-^Col1^+^CD14^+^ cells (**Figure 3A**) and were present at roughly half the proportion of activated CAFs (**Figure 3B**), which agreed with the scRNA-seq data (**Figure 2C**). Fibrocytes (EpCAM^-^CD31^-^CD11b^-^CD90^-^CD14^+^) from WiDr tumours were then isolated by fluorescence-activated cell sorting (FACS) for downstream qPCR analysis compared to sorted myeloid cells (EpCAM^-^CD31^-^CD11b^+^CD90^-^) and CAFs (EpCAM^-^CD31^-^CD11b^-^CD90^+^). We compared the scRNAseq data with qPCR data from sorted myeloid cells, fibrocytes, and CAFs for expression of *Ptprc* (CD45), *Itgam* (CD11b), *Cd14*, S100a8, *Col3a1*, and *Ly6c1* (**Figure 3C-D**), validating the relative abundance of these transcripts between the different cell types. We also validated CD45, CD11b, and CD14 protein expression in sorted myeloid cells, fibrocytes, and CAFs (**Figure 3E**) to confirm the identity of fibrocytes in these tumours. Our scRNA-seq data indicate that fibrocytes in telmisartan-treated tumours showed decreased abundance of *Col1a1*, *Col1a2*, and *Col3a1* transcripts and increased expression of MHC-class II antigen presentation genes *Cd74*, *H2-Ab1*, and *H2-Aa* compared to fibrocytes from control tumours (**Figure 3F**). We have therefore identified and validated a rare cell population of fibrocytes in WiDr tumours that respond to telmisartan treatment through a shift from fibrosis-associated genes to antigen presentation-related genes.

**Figure 3.**
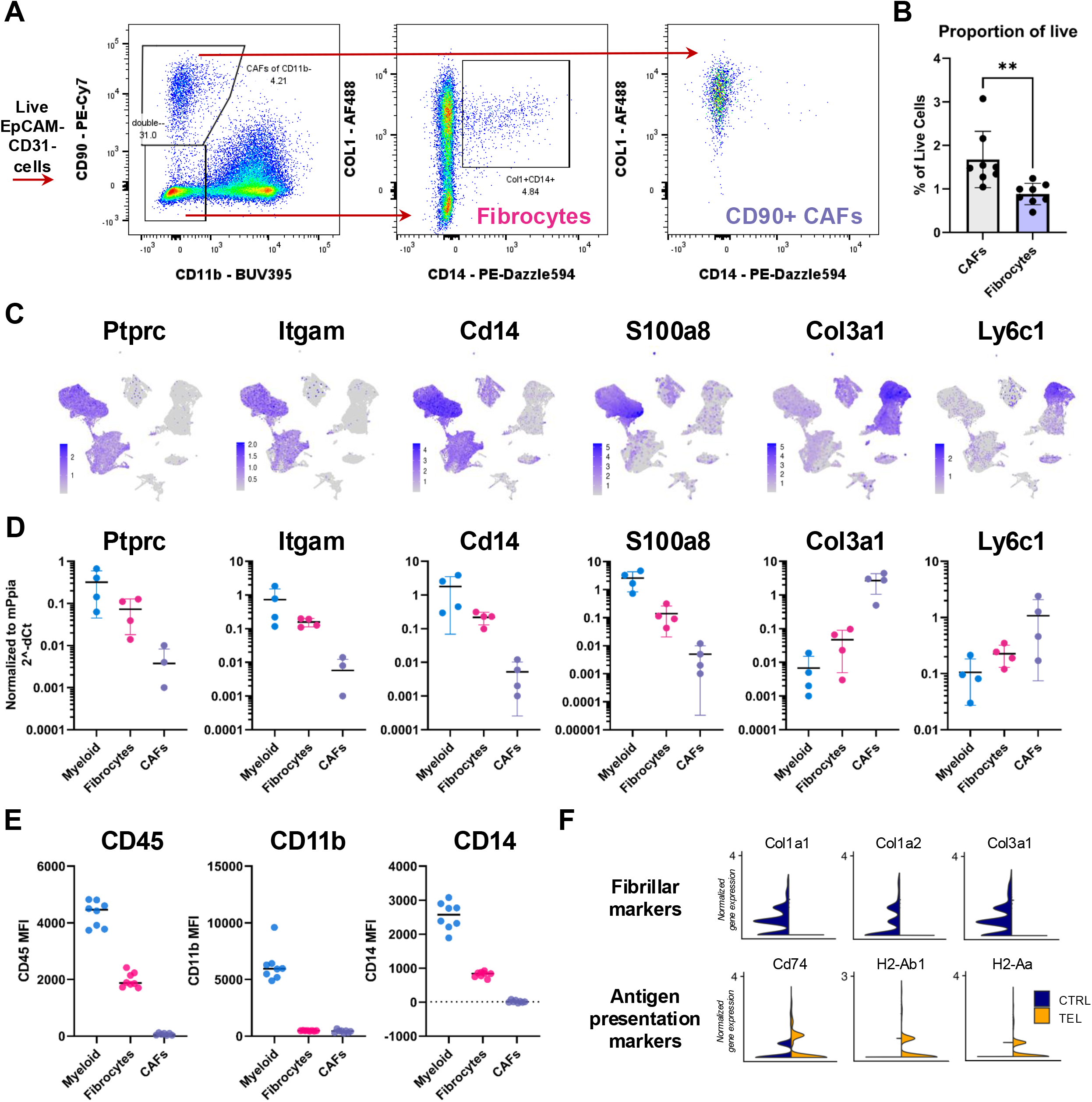
Fibrocytes characterized in WiDr tumours are affected by telmisartan treatment. **A)** Gating strategy for fibrocytes: live, EpCAM- and CD31-cells were gated into CD11b-CD90+ CAFs (CD90+ CAFs) or CD11b-CD90-cells which were further gated on Col1 and CD14 to define fibrocytes as CD11b-CD90-Col1+CD14+ fibrocytes. Col1 and CD14 marker expression was shown on CD90+ CAFs. **B)** The fibrocytes and CAF populations as a proportion of live. **C-D)** Gene expression of *Ptprc*, *Itgam*, *Cd14*, *S100a8*, *Col3a1*, and *Ly6c1* in **C)** scRNA-seq data, and **D)** qPCR conducted on FAC sorted myeloid, fibrocytes, or CAFs (sorted from similar gating strategy to **3A**, without Col1); qPCR values normalized to *Ppia* gene expression for the given cell type. **E)** Mean fluorescence intensity (MFI) of CD45, CD11b, and CD14 from flow cytometry of myeloid cells (EpCAM-, CD31-, CD11b+), fibrocytes, and CAFs. **F)** Fibrocyte expression of fibrillar markers (*Col1a1*, *Col1a2*, *Col3a1*) and antigen presentation markers (*Cd74*, *H2-Ab1*, *H2-Aa*) in control tumours (CTRL; navy) and telmisartan-treated tumours (TEL; yellow).

To further study the role of telmisartan on CAF subsets, we re-clustered the two CAF populations from the mouse UMAP object in Figure 2B (myCAFs and iCAFs, clusters 4 and 5) into a new UMAP projection (**Figure 4A**). At a resolution of 0.1, there were 5 clusters, one of which was found to be melanocytes which were removed from further analysis (**Supplementary Figure 3A-B).** The myCAF cluster (green) was defined by *Acta2*, *Tagln*, and *Thbs1*, while the iCAF cluster (orange) was identified by high levels of *Ly6c1*, *Dcn*, and *Efemp1* (**Figure 4B, C**). The proliferating CAFs or “prolifCAFs” (yellow) expressed moderate levels of both myCAF and iCAF markers, and relatively high levels of proliferation markers *Mki67*, *Cenpa*, and *Pclaf*. The undefined “nCAFs” (pink) had expression of both myCAF and iCAF markers (*Acta2, Tagln; Ly6c1, Dcn*), but also expressed dermal fibroblast markers *Apod*, *Cd9*, and *Wif1*. All clusters expressed the general CAF markers *Col1a1* and *Thy1* (**Figure 4C**). There was little change in CAF subset proportions between untreated and telmisartan-treated tumours (**Supplementary Figure 3C**).

**Figure 4.**
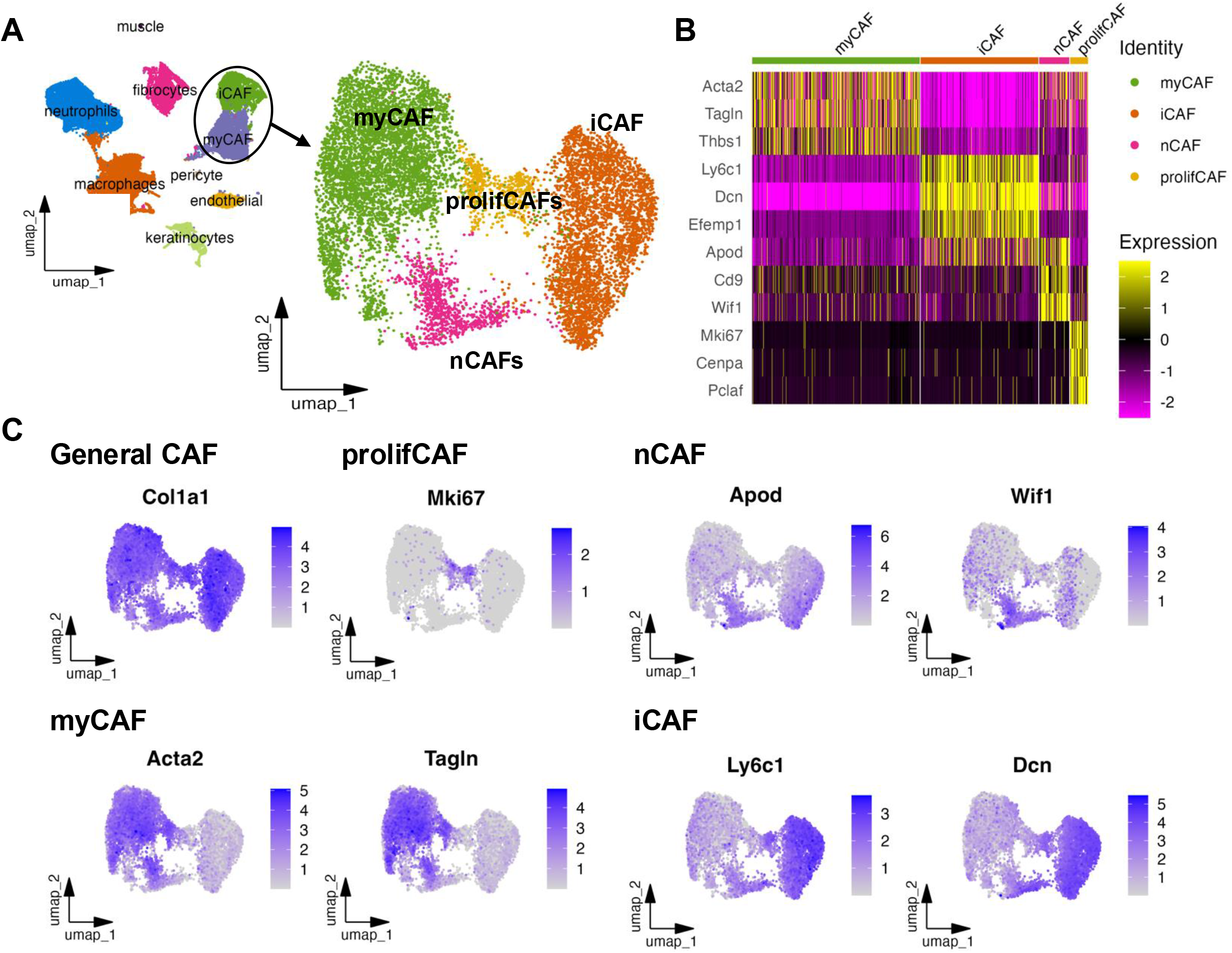
CAF populations re-clustered from mouse cells in scRNA-seq. **A)** Cells from the myCAF and iCAF clusters in the mouse UMAP projection were taken and created into a new UMAP projection of CAFs which were then clustered with Louvain clustering at a resolution of 0.1 to define four clusters, myCAFs, iCAFs, nCAFs, and prolifCAFs. **B)** Markers defining each cluster; myCAF (*Acta2*, *Tagln*, *Thbs1*), iCAF (*Ly6c1*, *Dcn*, *Efemp1*), nCAFs (*Apod*, *Cd9*, *Wif1*), and prolifCAFs (*Mki67*, *Cenpa*, *Pclaf*). **C)** Single marker UMAP projection of CAF markers.

To assess if telmisartan treatment caused phenotypic changes within each CAF cluster, differentially expressed gene (DEG) analysis followed by gene set enrichment analysis (GSEA) were conducted. Within the myCAFs, telmisartan treatment decreased gene signatures related to collagen deposition, including “ECM proteoglycans” (Reactome) and “ECM structural components” (GO Molecular Functions) (**Figure 5A**). Telmisartan treatment also increased myCAF gene signatures associated with immune modulation including “chemokine signaling” (WikiPathways) and “immunoregulatory interactions between a lymphoid and non-lymphoid cell” (Reactome) (**Figure 5A**). Transcriptional differences in the other three CAF populations were also present with telmisartan treatment. Telmisartan increased gene signatures related to epithelialization (“Keratinization”, Reactome) in iCAFs and nCAFs (**Figure 5B, C**), which is a common feature of the end stages of wound healing. We also observed signatures associated with MHC class II antigen presentation within the prolifCAFs of telmisartan-treated tumours (**Figure 5D**). Additionally, telmisartan induced a decrease in collagen/ECM-related gene signatures in iCAFs and nCAFs (**Figure 5B-C**), similar to our myCAF observations (**Figure 5A**). Overall, telmisartan decreased expression of ECM-related genes across several CAF subsets and increased gene profiles related to chemokine signaling, cell-cell interactions, keratinization, and antigen presentation.

**Figure 5.**
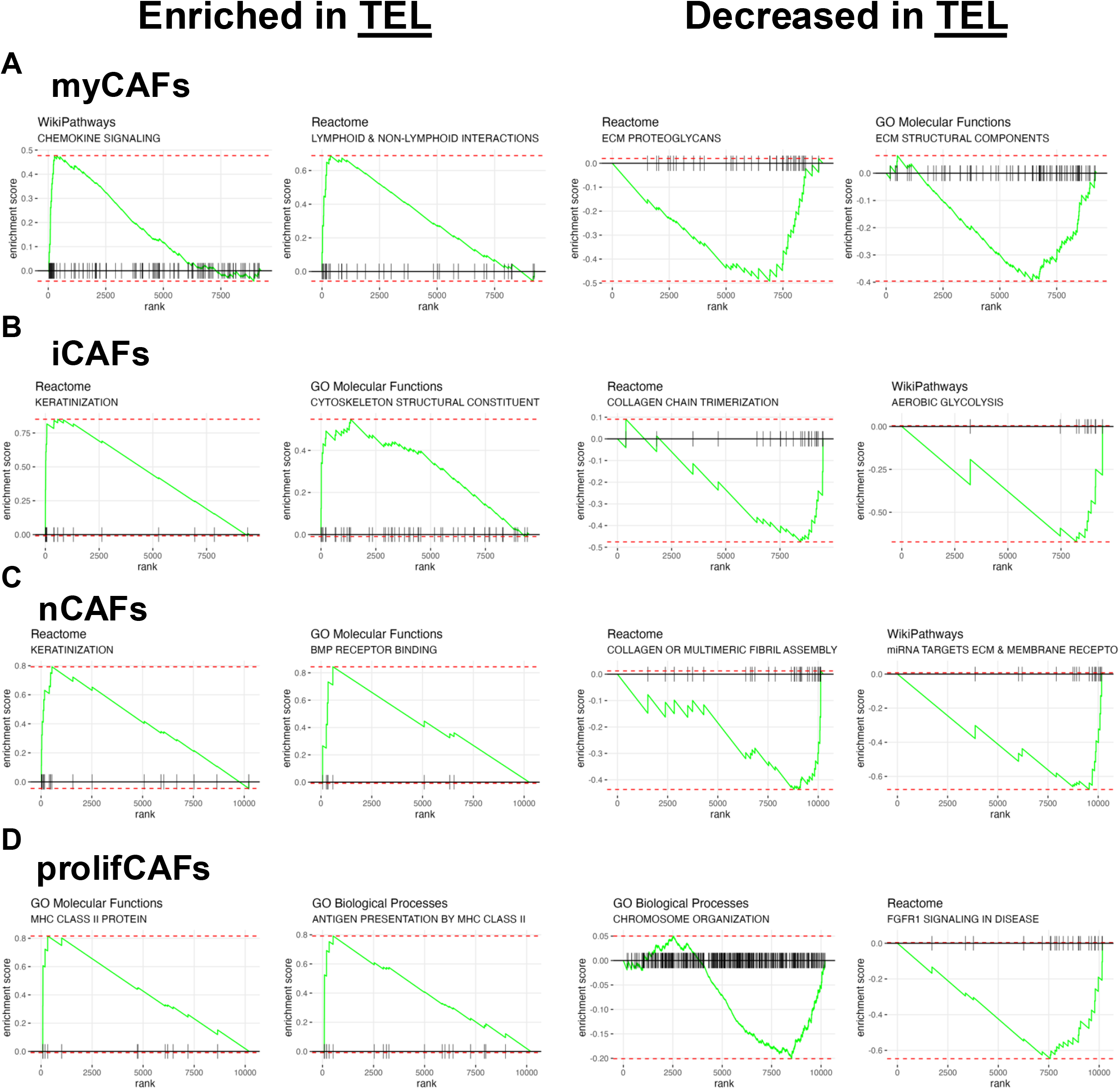
Changes in fibroblasts with telmisartan treatment at cluster resolution 0.1. Gene set enrichment analysis (GSEA) enrichment plots for: **A)** myCAFs: ECM proteoglycans (Reactome) and ECM structural constituent (GO molecular functions) enriched in CTRL, whereas chemokine signaling pathway (Wikipathways) and immunoregulatory interactions between a lymphoid and a non-lymphoid cell (Reactome) gene sets enriched in TEL myCAFs. **B)** iCAF: CTRL enriched for collagen chain trimerization (Reactome) and aerobic glycolysis (WikiPathways); TEL enriched for keratinization (Reactome), and structural components of cytoskeleton (GO Molecular Functions). **C)** nCAF: CTRL enriched for assembly of collagen fibrils and other multimeric structures (Reactome) and miRNA targets in ECM and membrane receptors (WikiPathways); TEL enriched for BMP receptor binding (GO Molecular Functions) and keratinization (Reactome). **D)** prolifCAFs: CTRL enriched for chromosome organization (GO Biological Processes) and signaling by FGFR1 in disease (Reactome); TEL enriched for MHC class II protein complex binding (GO Molecular Functions) and antigen processing and presentation of peptide or polysaccharide antigen via MHC class II (GO Biological Processes).

The telmisartan-induced phenotypic changes observed in CAFs through the GSEA data aligned with the observation of decreased αSMA expression in myCAFs by flow cytometry. We were curious whether the influence of telmisartan on CAFs could be further discriminated, particularly since the UMAP overlay comparing CAFs from control and telmisartan-treated tumours identified a distinct myCAF sub-population that is present only in tumours treated with telmisartan (**Supplementary Figure 3D**). We conducted higher resolution clustering of the scRNAseq data from WiDr tumour CAFs and found that a resolution of 0.5 in Louvain clustering identified 8 clusters: myCAF1, myCAF2, hypoxiCAF, telCAF, nCAF, prolifCAF, iCAF1, and iCAF2 (**Figure 6A**). The prolifCAF and nCAF clusters were maintained, while the larger myCAF and iCAF populations clustered into subgroups. Two new clusters were identified from the original myCAF population that displayed distinct gene signatures, including “hypoxiCAFs” that expressed transcripts associated with hypoxia and glucose metabolism gene signatures (**Supplementary Figure 3E**), and “telCAFs” that were enriched in transcripts for MHC class II antigen presentation including *H2-Ab1* and *Cd74* (**Supplementary Figure 3F**). The myCAF1 and myCAF2 subclusters, as well as the iCAF1 and iCAF2 subclusters, showed subtle transcriptomic differences (**Supplementary Figure 3G**), but overall shared the same genes that identified the larger myCAF and iCAF clusters identified with a 0.1 resolution, respectively. Within the higher resolution clustering of CAFs, we found that telmisartan increases the proportions of telCAFs and nCAFs while decreasing myCAF1 and myCAF2 subpopulations (**Figure 6B, C**). Both the telCAFs and nCAFs express lower levels of myCAF markers *Acta2* and *Col1a1* compared to myCAF1 and myCAF2 cells (**Figure 6D, E**). Additionally, the telCAFs and nCAFs express higher levels of iCAF markers *Ly6c1* and *Dcn* (**Supplementary Figure 3H-I**). Telmisartan therefore modifies the myCAF compartment of WiDr tumours to favor decreased *Acta2* and *Col1a1* expression.

**Figure 6.**
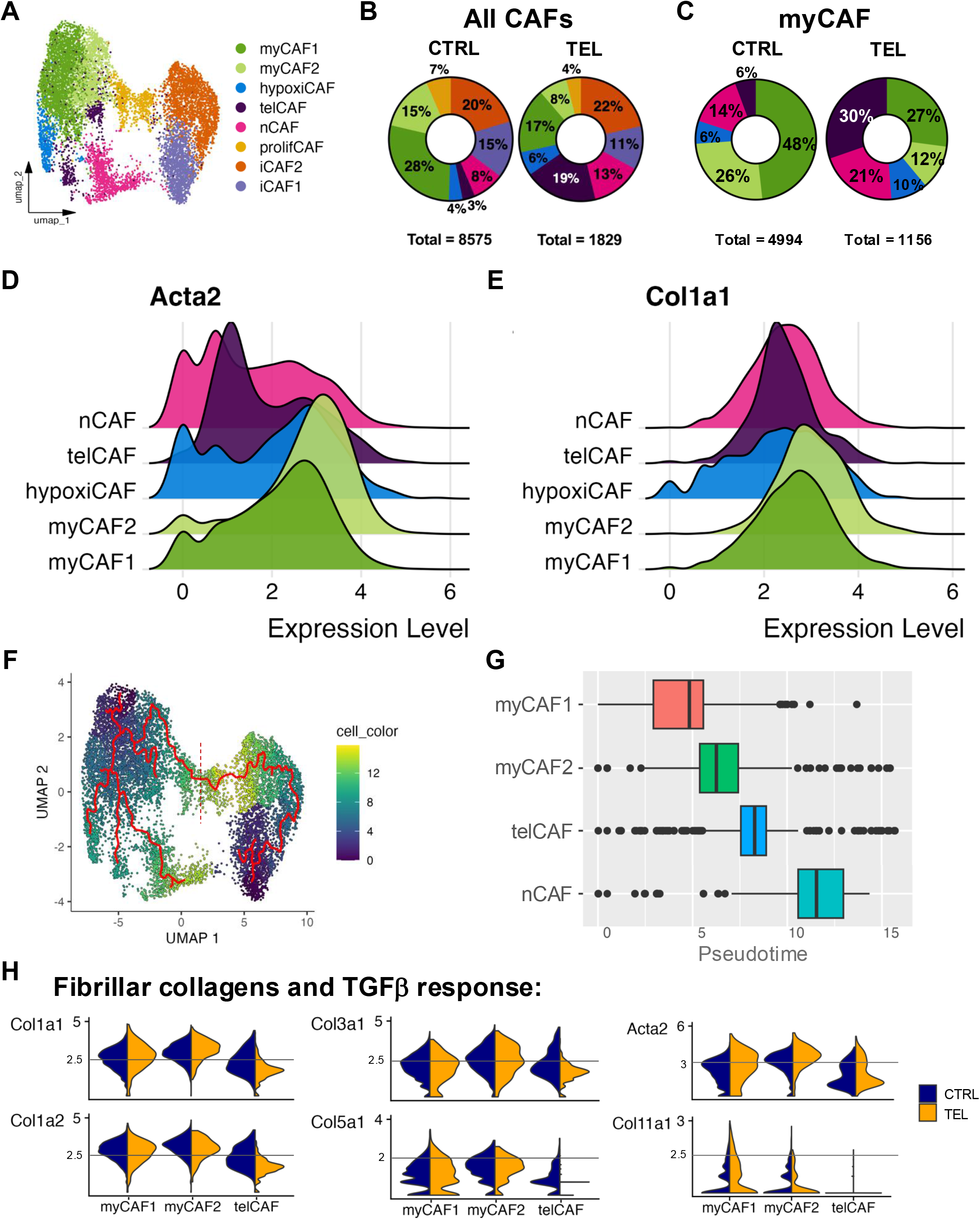
Higher resolution fibroblast clustering identifies myCAF subset affected by telmisartan treatment. **A)** Louvain clustering at a higher resolution of 0.5 identifies new clusters which are named as hypoxiCAFs, iCAF1, iCAF2, nCAF, myCAF1, myCAF2, prolifCAF, and telCAF. **B)** Proportion of each CAF cluster in control (CTRL) and telmisartan (TEL) treated tumours represented in pie charts. **C)** Changes to proportion of myCAF sub-populations (myCAF1, myCAF2, hypoxiCAF, and telCAFs) and nCAFs in CTRL and TEL CAFs. Ridgeplots of **D)** *Acta2* and **E)** *Col1a1* expression within myCAF and nCAF clusters. **F)** Pseudotime analysis setting one node each in iCAF1 and myCAF1 tract edges to show dynamic transcriptional changes across clusters. Dotted red line signifies the break in the prolifCAF cluster that distinguishes proliferating iCAFs vs myCAFs. **G)** Boxplot of pseudotime analysis quantification in myCAF1, myCAF2, and telCAFs. **H)** Violin plots to demonstrate expression levels of fibrillar collagen *Col1a1*, *Col1a2*, *Col3a1*, *Col5a1*, *Col5a3*, and *Acta2* within myCAF1, myCAF2, and telCAFs in CTRL (navy blue) and TEL (yellow). The Y-axis represents the relative expression of the indicated gene.

We hypothesized that telCAFs and nCAFs could be transcriptionally derived from the more myofibroblast-like myCAF1 or myCAF2 clusters. Pseudotime analysis produced a pseudotime track showing that the telCAFs, hypoxiCAFs, and nCAFs all derived from the myCAF1 cluster (**Figure 6F**). The telCAF population showed a pseudotime track that was distinct from the nCAFs. Additionally, the telCAFs and nCAFs are transcriptionally distinct from the myCAF1 and myCAF2 populations as their relative pseudotime number is higher (**Figure 6G**). Violin plots of myCAF1, myCAF2, and telCAFs from control (yellow) or telmisartan-treated (navy blue) tumours showed that expression of fibrotic collagens (*Col1a1, Col1a2, Col3a1, Col5a1,* and *Col11a1*) was lower in telCAFs compared to myCAF1 and myCAF2 populations. Further, telmisartan treatment decreased the frequency and number of each of these collagen transcripts within the telCAF populations with minimal effects on these genes within the myCAF1 and myCAF2 populations (**Figure 6H**). Our data demonstrate that telmisartan alters the transcriptome of myCAFs toward a less fibrotic phenotype, providing a mechanistic explanation for decreased collagen deposition observed in WiDr tumours treated with^17^.

## Discussion

In this work, we identify multiple CAF subsets and fibrocytes in a solid tumour xenograft model and we show that telmisartan induces transcriptomic changes in fibrocytes and myCAFs that result in reduced expression of several fibrosis-related genes. These data are consistent with our previously published work showing that telmisartan decreases collagen deposition in WiDr tumours^17^, providing further support for the use of telmisartan to manipulate cellular components of the solid tumour microenvironment. Our data also indicate that telmisartan increases the relative expression of transcripts associated with antigen presentation in the fibrocyte and myCAF populations.

In humans, fibrocytes arise from circulating CD14+ monocytes that infiltrate wounds and become fibroblast-like. Fibrocytes may express myeloid markers (S100A8/9, PTPRC), antigen presenting markers (CD74, MHC Class II genes) and fibrillar collagen markers (COL1A1, COL3A1, ACTA2)^6,7,10,18^. In mice, fibrocytes are not as well-characterized but similarly to human fibrocytes, express both myeloid cell and fibroblast-associated genes. In both humans and mice, circulating fibrocytes have been shown to lose expression of hematological markers including CD45 and CD11b when the cells enter tissues^10^. In our study we demonstrate the presence of these rare fibrocytes in a human tumour model through scRNA-seq analyses and a newly developed flow cytometry protocol. Fibrocytes have been previously difficult to study in other mouse models and human samples, so our work has created a new framework for studying fibrocytes in cancer and other disease models. Additionally, we show that telmisartan decreases expression of collagen transcripts in fibrocytes despite minimal effects on myeloid cell transcriptomes in these tumours. Combined with the telmisartan-induced decrease in collagen expression by myCAFs, these data indicate that telmisartan reduces fibrosis-associated genes in multiple cell types that may contribute to fibrosis, regardless of the cell type of origin. Future *ex vivo* work with cultured fibrocytes would help to understand how these cells respond to telmisartan (or other ARBs) and elucidate the contribution of fibrocytes to the fibrotic extracellular matrix found in some tumour types.

We observed significant CAF heterogeneity in WiDr tumours as observed through higher resolution analyses of our scRNA-seq data, including two distinct myCAF subclusters (telCAF and nCAF) that proportionally increased in telmisartan-treated tumours. These telCAFs and nCAFs had reduced expression of myCAF-related genes such as *Acta2* and *Col1a1*, and had higher iCAF-related gene expression (*Ly6c1* and *Dcn)*. These two subclusters were transcriptionally distinct from each other and evolved separately from the myCAF1 and myCAF2 subclusters in our pseudotime analysis. We speculate this may be due to differences in exposure of myCAFs to cytokines within the tumour microenvironment. Fibroblasts exposed to IL-1 typically transition to an iCAF phenotype, while fibroblast exposed to TGFβ transition toward a myCAF phenotype. While CAFs in a complex microenvironment can be exposed to both IL-1 and TGFβ, these cells tend to maintain more of a myCAF phenotype since TGFβ, signaling outcompetes IL-1 signalling. The nCAFs in our analyses may represent a CAF population that is responding to competing IL-1 and TGFβ in the environment, whereas telCAFs may represent the direct effect of telmisartan on myCAFs. Further experiments exposing myCAFs and iCAFs to telmisartan *ex vivo* would help to elucidate how telmisartan directly impacts CAF phenotype and how telmisartan influences the response of different CAF populations to cytokines.

Currently, there are no FDA approved therapies that specifically target myCAFs in solid tumours to act as anti-fibrotics in cancer. However there are several putative anti-fibrotics for cancer treatment being studied for cancer treatment, including the pan-lysyl oxidase inhibitor PXS-5505 (Pharmaxis) to inhibit ECM crosslinking and reduce tumour stiffness in PDAC^19^ and systemic sclerosis^20^, which is currently in clinical phase 2 trials for myelofibrosis^21^. Other therapeutic strategies including targeting fibroblast activating protein (FAP) on CAFs through CAR-T cells^22,23^, FAP-targeting antibodies^24^, inhibiting TGFβ / TGFβ binding receptors^30^ through neutralizing antibodies ^25–28^, or small molecule inhibitors such as galunisertib^29–33^. Unfortunately, many of the clinical trials associated with these therapies failed due to unacceptable side effects, little-to-no response to the medication, and/or poor patient selection for the studies. There is an important unfilled need for drugs that can specifically and safely target fibrosis-inducing myCAFs to modify the solid tumour microenvironment and improve tumour control in conjunction with existing cytotoxic therapies.

Multiple retrospective clinical studies have shown that ARB use is correlated with improved survival outcomes in various cancer types^34^ including advanced colorectal cancer^35,36^, metastatic kidney cancer^37^, pancreatic cancer^38,39^, lung cancer^40^, and our recent study in oropharyngeal cancer^41^. The relatively low proportion of cancer patients taking ARBs for hypertension complicates potential retrospective analyses assessing treatment responses from patients taking any particular ARB. The ARB losartan has been tested in a phase II clinical trial as an adjuvant therapy to FOLFIRINOX in pancreatic cancer patients^15^. This study demonstrated that losartan improved patient response to chemotherapy and subsequent proton radiotherapy, allowing a proportion of tumours to be surgically resected. We have previously shown that telmisartan induces greater inhibition of collagen deposition than losartan in a pre-clinical tumour model^17^, suggesting that telmisartan may prove more beneficial than losartan for reducing ECM deposition and improving tumour perfusion. Indeed, telmisartan exhibits a higher binding affinity for the angiotensin II receptor and has better bioavailability and increased circulation half-life than losartan^42,43^. There is currently one clinical trial testing telmisartan as a prostate cancer treatment neoadjuvant in the context of DNA repair^44,45^, although the use of telmisartan for decreasing tumoural fibrosis in cancer patients has not yet been tested.

In this work, we show that telmisartan modifies the cellular composition of a solid tumour model, altering the proportions of myeloid cells, CAFs, and fibrocytes. Telmisartan also modified the transcriptomes of multiple CAF subsets and fibrocytes, including decreasing the expression of extracellular matrix components by myCAFs and fibrocytes. These data indicate that telmisartan preferentially reduces the pro-fibrotic phenotypes of collagen-producing cells in solid tumours, and provides further support for testing telmisartan as a therapeutic adjuvant to decrease tumour fibrosis and improve cancer therapy efficacy.

## Methods

### Mouse tumour model

Immunocompromised NRG (NOD-Rag1null-Il2rg) mice were obtained from Jackson Laboratories (JAX) and bred in-house in the Animal Resource Centre at the BC Cancer Research Centre. Mice were implanted with a human cell line subcutaneously into the back left flank for flow cytometry and immunofluorescence (IF) microscopy or in contralateral flanks for scRNA-seq analyses. Tumours were collected when they reached around 300 – 500 mm^3^ based on length (L) and width (W) measurements obtained using calipers where tumour volume = (V x W^2^)/2

### Human cell line

The WiDr human colon adenocarcinoma cell line (CCL-218), was obtained from American Type Culture Collection (ATCC). WiDr cells were grown in minimum essential media (MEM) supplemented with 10% fetal bovine serum (FBS) (Gibco, 12483-020, origin: CanadaCells in exponential growth phase were trypsinized and rinsed with phosphate-buffered saline (PBS) two times prior to counting with a Countess 3 FL automated cell counter. Cells were resuspended in a 50% mix of serum-free media and PBS prior to being subcutaneously implanted into mice at a volume of 0.1 mL in a concentration of 10 million cells/mL.

### Tumour cell processing for flow cytometry

Mice were euthanized and tumours were removed when they were approximately 300 – 500 mm^3^. Tumours were minced with scalpels then added to an enzyme cocktail of 1 mg/mL of collagenase I and collagenase II each (Gibco, 17100017 & 17101015) in 3 mL of serum-free RPMI-1640 media. The minced tumours and enzyme cocktail were placed in a 37°C shaking incubator for 45 minutes. Afterwards, 1mg/mL DNAse (Sigma-Aldrich, DN25-1G) and 5 mL media with 10% FBS were added to the samples before pressing the contents with the rubber end of a syringe stopper through a 100 μm bucket filter. The resultant samples were centrifuged at 150 x g and filtered again through a 40 μm bucket filter. If there were visible clumps of cells leftover, samples were again filtered through a filter-top 5 mL tube. Cells were counted prior to adding viability dye and extracellular antibodies for 30 minutes, washed with HFN buffer (<u>H</u>ank’s neutral salt solution [Stemcell, 37150], 2% <u>F</u>BS, and 0.1% <u>N</u>aN_3_ sodium azide), then fixed and permeabilized using the FoxP3 / transcription factor staining buffer set (Invitrogen, 00-5523-00). For flow cytometry analysis, cells were fixed and permeabilized, intracellular antibodies were added overnight, then washed off, and the cells were resuspended in HFN for flow cytometry analysis.

### Tumour processing for scRNA-seq

Single-cell RNA-sequencing was done on two batches of 8 mice/group (control, “CTRL” or telmisartan, “TEL”), with 2 tumours/mouse (on each flank). Each group of 8 mice was considered one biological replicate in the final dataset. Whole tumours from both flanks were removed, minced, then added into an enzyme cocktail of 250 μL of 5 U/mL of neutral dispase (Worthington, LS02100) and 1 mg/mL of collagenase I and II each in 3 mL PBS then placed in a 37°C shaking incubator for 30 minutes. Afterwards, 1 mg/mL of DNAse then RPMI-1640 media with 10% FBS were added to saturate proteolytic enzymes in each sample. The samples were pressed through a 40 μm bucket filter using a syringe plunger then centrifuged at 150 x g. Cells were resuspended in PBS, and refiltered through a filtertop 5 mL tube (Falcon, 0877123). Samples were pooled by condition (CTRL or TEL) prior to red blood cell lysis with 9 mL of ACK lysis buffer (Gibco, A1049201) for 5 min at room temperature. After centrifuging at 150 x g, debris was removed with a debris removal kit (Miltenyi Biotec, 130-109-398) prior to treatment with human EpCAM-PE (BioLegend, 324206, Clone 9C4).

Magnetic bead PE-positive selection (EasySep; STEMCELL, 17684) was used to remove cancer cells. The flow through from the selection was then washed once in scRNA-seq buffer (Ca^2+^ and Mg2^+^ free PBS + 0.04% BSA) then reconstituted up to 200 μL of the scRNA-seq buffer and counting on a hemocytometer with trypan blue to ensure high viability and no visible debris.

### Flow cytometry staining

Flow cytometry was conducted on the single cell suspensions isolated from tumours. Prior to flow cytometry analysis, eBioscience fixable viability dye eFluor780 (Invitrogen, 65-0865-14) was added at a concentration of 0.1 μL / million cells for 30 min on ice. Cells were rinsed in HFN, placed in Fc block (anti-mouse CD16/32 Antibody, BioLegend, 101320) at 1 μL / sample for 15 min, then extracellular antibodies were added directly following Fc block at a concentration of 0.2 μL/ test unless otherwise specified. The WiDr extracellular antibodies panel had the following antibodies: anti-mouse CD45 BUV737 (0.1 μL/test, Invitrogen, 367-0451-82, Clone 30-F11), anti-mouse CD31 PE (BioLegend, 102408, Clone 390), anti-human EpCAM PE (0.6 μL/test, BioLegend, 324206, Clone 9C4), anti-mouse CD90 BV650 (BioLegend, 140318, Clone 53-2.1), anti-mouse PDPN PE-Cy7 (BioLegend, 127412, Clone 8.1.1), anti-mouse Ly6C PerCP-Cy5.5 (BioLegend, 128012, Clone HK1.4). Intracellular staining was done for anti-mouse αSMA eFluor 660 (Invitrogen, 50-9760-82, Clone 1A4) and collagen 1 (0.1 μL/test, Kerafast, ENH018-FP, Clone LF-68). Flow cytometry was conducted on the BD FACSymphony^TM^ A5 Cell Analyzer. Data were captured and analyzed on FlowJo Version 10 software, where compensation, gating, and quantitative data analyses were conducted. GraphPad Prism version 9 was used for statistical testing and graph creation.

### Tumour processing for cell sorting then RNA extraction

Tumours were processed as they were for scRNA-seq, with the PE-selection kit on EpCAM-PE stained cells (using the negatively selected flow-through). Once the flow through was obtained, cells were then processed similarly to the flow cytometry protocol for extracellular antibody staining. Cells were then run through the BD FACSAria™ Fusion Flow Cytometer for four-way cell sorting, to obtain myeloid cells (Live cells, EpCAM^-^, CD11b^+^), fibrocytes (Live cells, EpCAM^-^, CD11b^-^CD90^-^, CD14^+^), myCAFs (Live cells, EpCAM^-^, CD11b^-^CD90^+^, CD29^hi^, Ly6C^lo-mid^), and iCAFs (Live cells, EpCAM^-^, CD11b^-^CD90^+^, CD29^mid^, Ly6C^hi^). Cells were sorted into 150 μL of RLT lysis buffer from the RNeasy Mini Kit (QIAGEN, 74104). 200 μL of RLT lysis buffer was added to samples prior to being processed with the RNeasy kit. RNA was measured, and cDNA was synthesized with the High-Capacity RNA-to-cDNA™ Kit (Applied Biosystems, 4387406). If cells were referred to as CAFs, with no myCAF or iCAF designation, cDNA from myCAF and iCAF samples were pooled for qPCR for ease of reference to fibrocytes and immune cells.

### Quantitative Polymerase Chain Reaction (qPCR)

Primers for qPCR were designed and validated for mouse Ptprc, Itgam, Cd14, S100a8, Col3a1, and Ly6c1 genes. Primer stocks of 5 μM were used with Fast SYBR green (Applied Biosystems, 4385612). Primers are as follows:

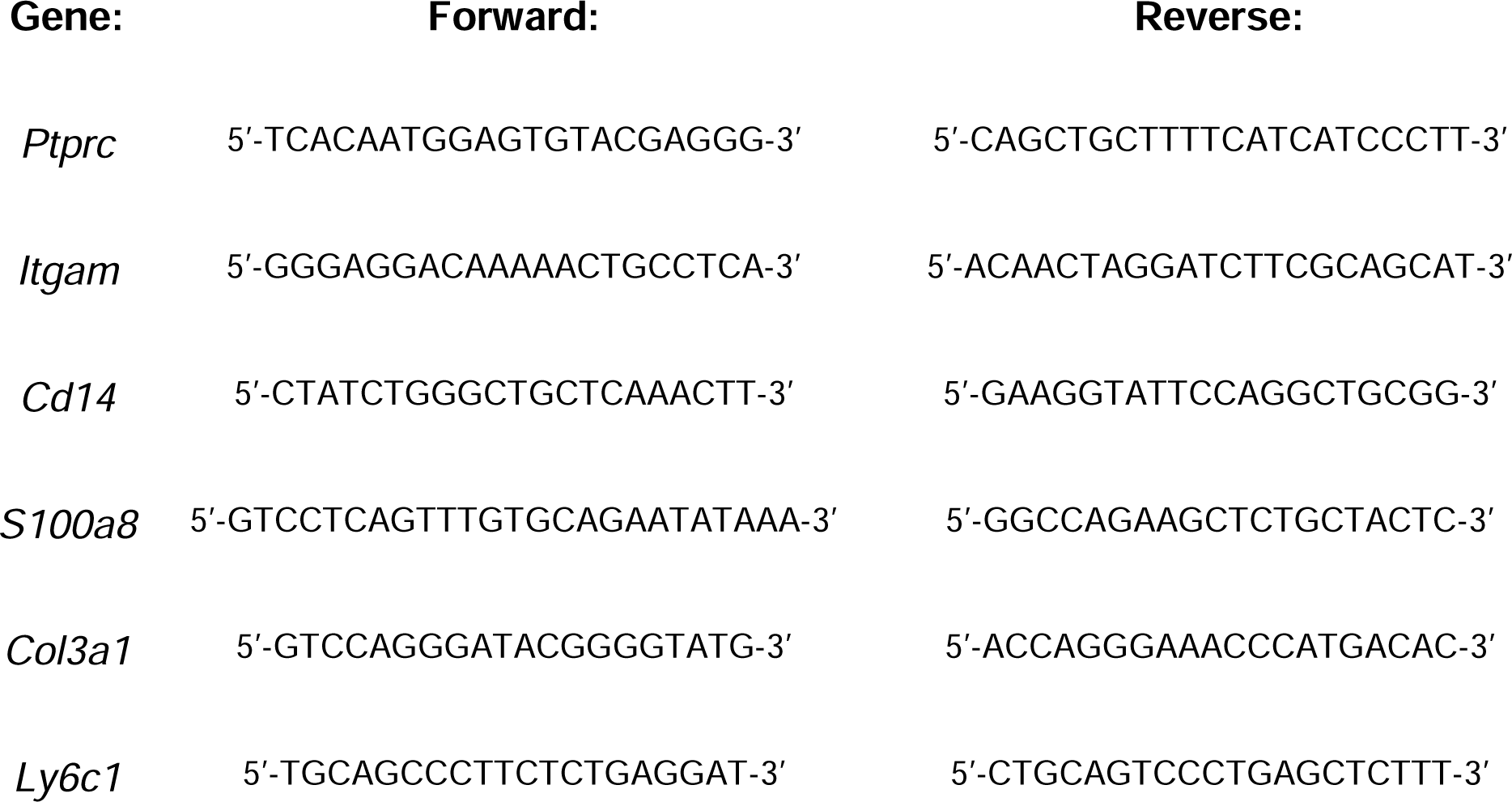

## scRNA-seq

### Sample running and quality control

Cells were processed for scRNA-sequencing using the 10x Chromium 3’ library preparation and sequencing at 100 million reads / sample at the University of British Columbia Bioinformatics School of Biomedical Engineering Sequencing Core. Cells were analyzed through the cellranger-6.0.1 pipeline and mapped to the GRCm38^46^ mouse genome prior to export. Data were then processed in R studio^47^ on R version 4.4.3^48^ using Seurat version 5^49,50^. Cells were included based on the following criteria: defined as singlets using DoubletFinder^51^, number of unique molecular identifiers (UMI) >= 500, at least 250 transcripts (nFeature_RNA) and less than 8000 transcripts, cells had a novelty score > 0.8 with novelty score defined as: log10(nFeatureRNA) / log10(UMI), and a mitochondrial fraction <20%. Genes were only included if they were expressed in 10 or more cells. Each sample was normalized with SCTransform prior to being integrated with SelectIntegrationFeatures and all samples merged back into one dataset. Principal component analyses showed that there was minimal batch effect between samples. Once cells passed this pipeline of quality control, FindNeighbors (dims 1:40) and FindClusters were both conducted prior to a Universal Manifold Approximation and Projection (UMAP) being created for further downstream analysis. Louvain clustering was conducted to define clusters within the mouse or fibroblast UMAPs. Cluster resolution is noted in text.

### Cluster definition, differentially expressed genes (DEG) and GSEA analysis

R studio was used for data processing, including data.table^52^, dplyr^53^, magrittr^54^, qs^55^, R.utils^56^, and tidyverse^57^ to aid with coding and software running facility; ggplot2^58^, cowplot^59^, patchwork^60^, viridis^61^, and scCustomize^62^ for data visualization; multtest^63^, metap^64^, and presto^65^ for statistical analysis. To define clusters FindAllMarkers with the MAST test for differential genes was conducted, and graphs were created to best visualize cluster identities based on known markers. SingleR^66^ was initially conducted to help identify and validate clusters to define certain cell types. To better define subclusters or to observe phenotypic differences within a cluster between CTRL and TEL, FindMarkers was conducted between clusters of interest to create a log2FC ranked gene list that could be used for subsequent gene set enrichment analysis (GSEA). GSEA was conducted on a descending list of Log2FC genes defined from the above FindMarkers function for a given cluster. Then, fgsea^67^ was used to conduct GSEA within various msigdbr^68–70^ collections to define the top enriched pathways. The adjusted p-value and normalized enrichment score (NES) were used for these pathway analyses. The msigdbr collections used were: CGP, CP (Biocarta, Reactome, Wikipathways), GTRD, GO Molecular Functions, GO Biological Processes, Hallmarks, and CellTypes (collection 8). Specific pathways of interest were plotted using plotEnrichment and leading-edge genes were also used to identify if pathways of interest were properly representing the differences in gene enrichment.

### Pseudotime analysis

Pseudotime analysis was conducted using the Monocle3 package v1.3.7 ^71–74^ and SeuratWrappers^75^ in Seurat on the CAF population shown in Figure 4. Pseudotime was used to define how clusters were defined branching off of myCAF and iCAF populations. Thus, root nodes were defined as the clusters “myCAF1” and “iCAF1” based on cluster resolution 0.5.

## DATA AVAILABILITY STATEMENT

R scripts used to analyze these data can be provided upon request.

## ANIMAL ETHICS

Ethics for this work was approved through the animal research protocol A21-0266 and A25-0214 by the UBC Animal Care Committee in compliance with the Canadian Council on Animal Care.

## CRediT AUTHORSHIP CONTRIBUTION STATEMENT

**Che-Min Lee**: Conceptualization, Data Curation, Formal analysis, Investigation, Methodology, Software, Writing – original draft, Visualization. **Nikita Telkar**: Software, Writing – review & editing. **Brennan J. Wadsworth**: Investigation, Methodology. **Rachel A. Cederberg**: Methodology. **Rocky Shi**: Software. **Lisa Zhan:** Investigation. **Prabhreet Sekhon:** Investigation. **Meredith A. Clark:** Investigation. **Kiersten N. Thomas:** Investigation. **Michael G.S. Hall**: Investigation. **Wan L Lam**: Supervision. **Kevin L. Bennewith**: Conceptualization, Funding Acquisition, Resources, Supervision, Writing – review & editing.

## ACKNOWLEDGEMENTS

We would like to thank the UBC Sequencing and Bioinformatics Consortium, namely facility manager Tara Stach and bioinformaticians Bernie Zhao for initial CellRanger data processing and Christine Yanta for quality control, processing, and helping visualization in Seurat. We would also like to thank the staff from the Animal Resource Centre and Flow Cytometry core at BC Cancer Research Centre for their assistance and expertise. We would like to thank Jasmin Wachter for advice on scRNA-seq sample preparation and Quincy Collins and Dr. Michael Underhill for helpful discussion on scRNA-seq analysis.

## SUPPLEMENTARY INFORMATION

Supplementary information (pdf) attached.

---

**Supplementary Figure 1.**
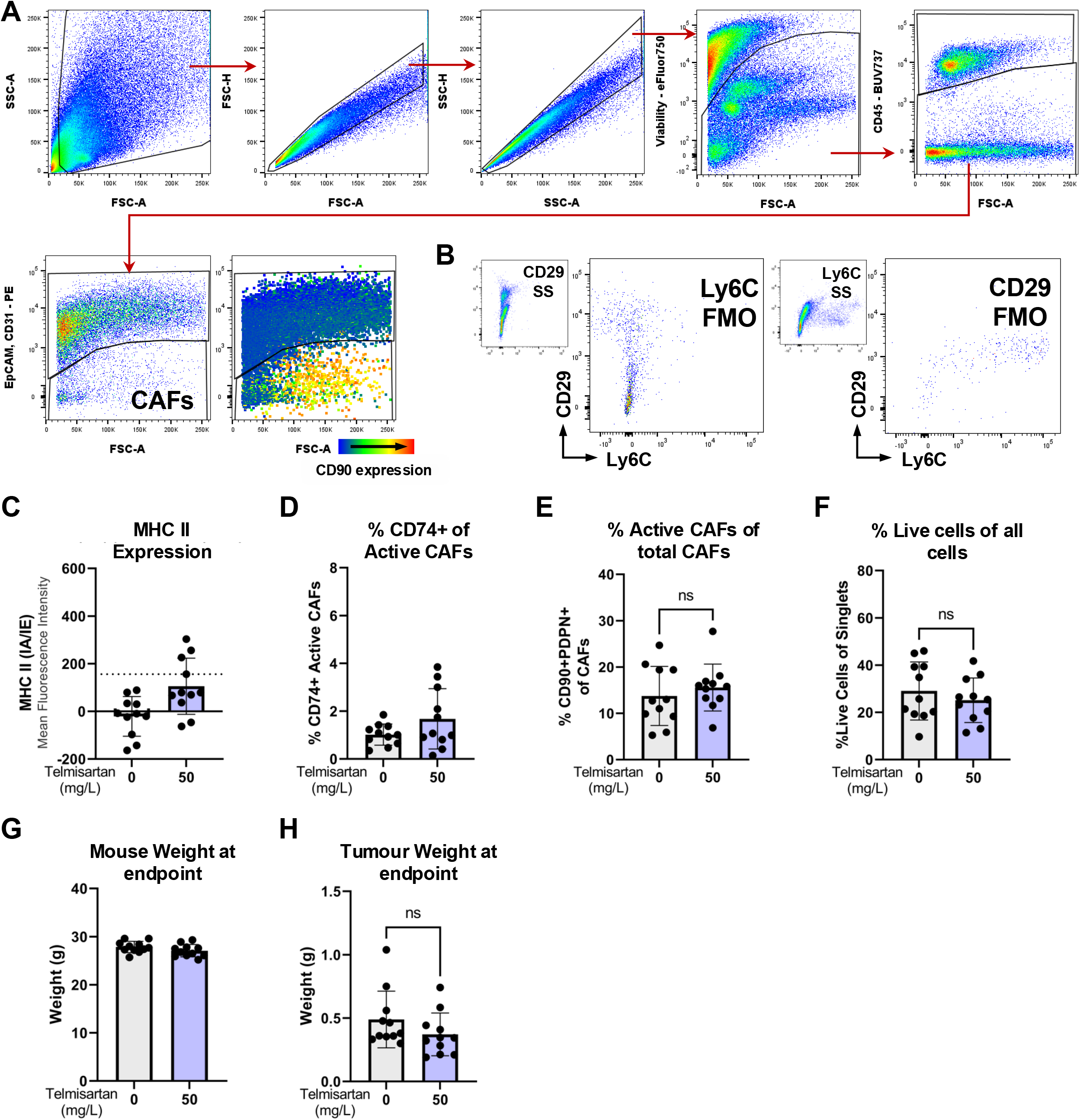
Flow gating strategies and experimental controls. **A)** Gating strategy for to find live, singlet, CD45-, EpCAM-, CD31- CAF populations in WiDr tumours. **B)** Fluorescence minus one (FMO) controls for Ly6C and CD29. **C)** MHC II (IA/IE) mean fluorescence intensity in PDPN+CD90+ CAFs of WiDr tumours +/− telmisartan treatment. Dotted line represents the FMO control. **D)** Proportion of CD74+ CAFs within active PDPN+CD90+ CAFs. **E)** Active CAFs as a proportion of total CAFs. **F)** Total live cells (negative for viability dye) of singlets (all cells). **G)** Mouse weights at experimental endpoint in control (grey) or telmisartan (blue) treated tumours. **H)** WiDr tumour weights at experimental endpoint.

**Supplementary Figure 2.**
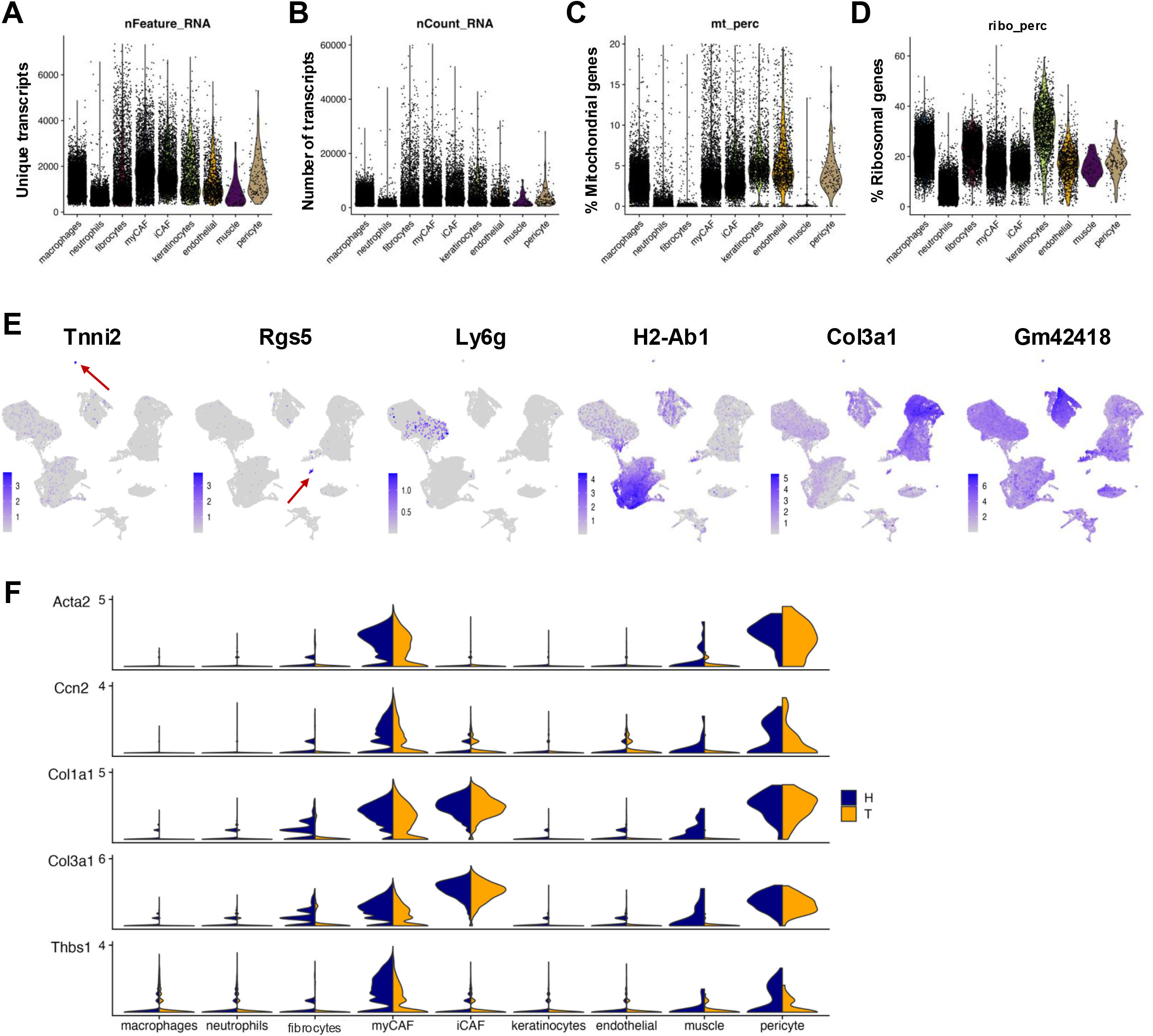
Quality control metrics, extra marker expression, and fibrocyte gating strategy and markers in WiDr mouse cells. **A-D)** Quality control metrics of each cluster in mouse cells of WiDr tumours. **A)** Total number of unique transcripts in each cluster, column called “ nFeatures_RNA” in dataset. **B)** Number of total transcripts, called “nCount_RNA” in dataset. **C)** Percentage of mitochondrial gene (gene name starting with ‘mt-”) fraction, “mt_perc” in dataset. **D)** Percentage of ribosomal gene (gene name starting with “Rp”) fraction, “percent.ribo” in dataset. **E)** Additional clustering markers *Tnni2* (muscle), *Rgs5* (pericytes), *Ly6g* (neutrophils/granulocytes), *H2-Ab1* (antigen presentation; macrophages, fibrocytes), *Col3a1* (fibroblasts, fibrocytes), and *Gm42418* (all, highest in fibrocyte cluster). **F)** Fibrillar collagens (Col1a1, Col3a1), myofibroblast marker Acta2, TGFb-related genes thrombospondin-1 (Thbs1), and TGFb downstream target connective tissue growth factor (Ccn2) expression across all cells in WiDr tumours.

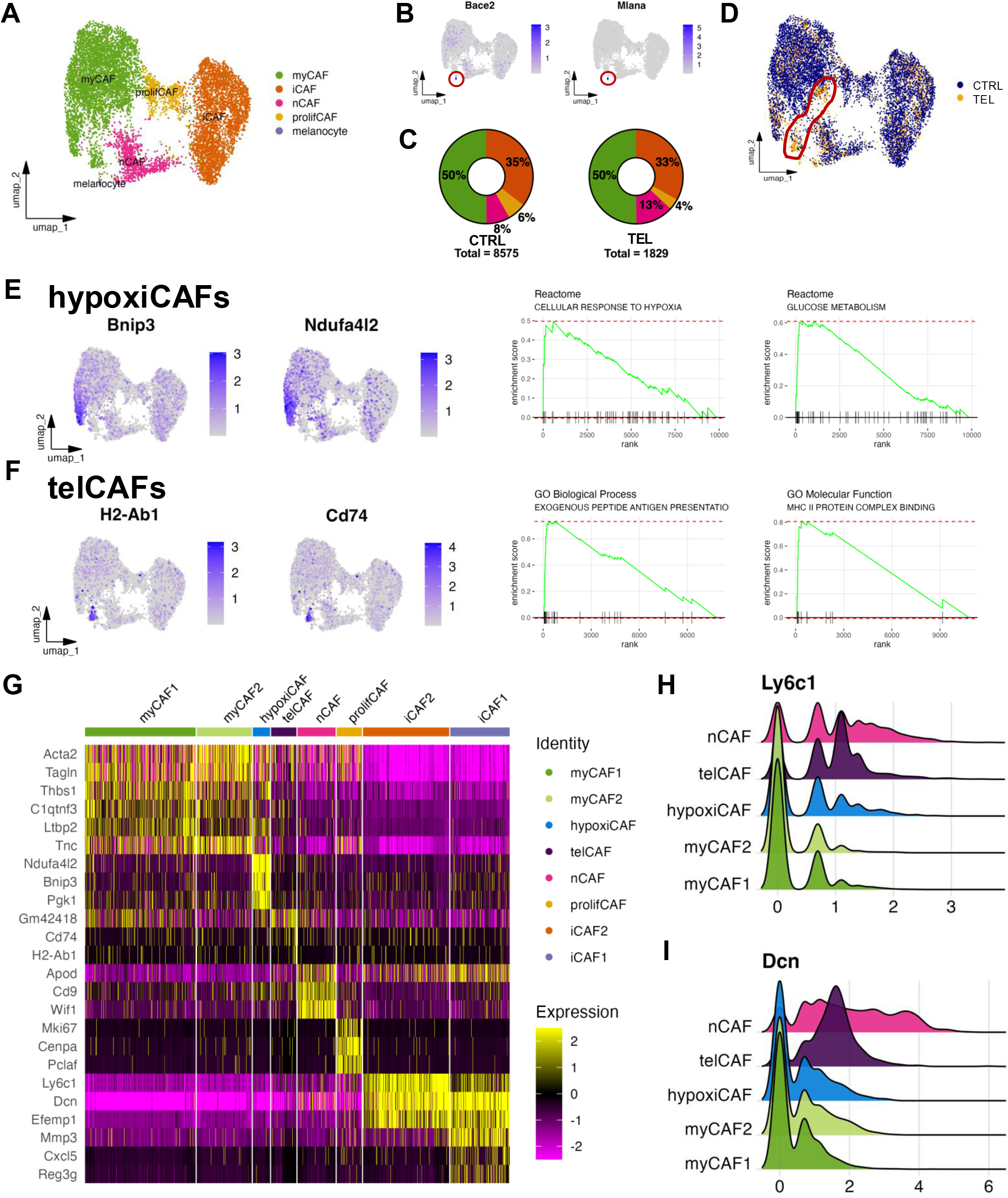

## References

1. Kalluri, R. The biology and function of fibroblasts in cancer. Nat. Rev. Cancer 16, 582–598 (2016).

2. Paraiso, K. H. T. & Smalley, K. S. M. Fibroblast-mediated drug resistance in cancer. Biochem. Pharmacol. 85, 1033–1041 (2013).

3. Farmer, P. et al. A stroma-related gene signature predicts resistance to neoadjuvant chemotherapy in breast cancer. Nat. Med. 15, 68–74 (2009).

4. Hanley, C. J. et al. Targeting the Myofibroblastic Cancer-Associated Fibroblast Phenotype Through Inhibition of NOX4. J. Natl. Cancer Inst. 110, 109–120 (2018).

5. Li, B. et al. Cell-type deconvolution analysis identifies cancer-associated myofibroblast component as a poor prognostic factor in multiple cancer types. Oncogene 2021 4028 40, 4686–4694 (2021).

6. Roife, D., Fleming, J. B. & Gomer, R. H. Fibrocytes in the Tumor Microenvironment. Adv. Exp. Med. Biol. 1224, 79–85 (2020).

7. Weigert, A. et al. Fibrocytes boost tumor-supportive phenotypic switches in the lung cancer niche via the endothelin system. Nat. Commun. 13, 6078 (2022).

8. Öhlund, D. et al. Distinct populations of inflammatory fibroblasts and myofibroblasts in pancreatic cancer. J. Exp. Med. 214, 579–596 (2017).

9. Bernard, V. et al. Single-Cell Transcriptomics of Pancreatic Cancer Precursors Demonstrates Epithelial and Microenvironmental Heterogeneity as an Early Event in Neoplastic Progression. Clin. Cancer Res. 25, 2194–2205 (2019).

10. Suga, H. et al. Tracking the elusive fibrocyte: Identification and characterization of collagen producing hematopoietic lineage cells during murine wound healing. Stem Cells Dayt. Ohio 32, 1347–1360 (2014).

11. Cohen, C. et al. WNT-dependent interaction between inflammatory fibroblasts and FOLR2+ macrophages promotes fibrosis in chronic kidney disease. Nat. Commun. 2024 151 15, 1–23 (2024).

12. Timperi, E. et al. Lipid-Associated Macrophages Are Induced by Cancer-Associated Fibroblasts and Mediate Immune Suppression in Breast Cancer. Cancer Res. 82, 3291–3306 (2022).

13. Piwocka, O., Piotrowski, I., Suchorska, W. M. & Kulcenty, K. Dynamic interactions in the tumor niche: how the cross-talk between CAFs and the tumor microenvironment impacts resistance to therapy. Front. Mol. Biosci. 11, (2024).

14. Chauhan, V. P. et al. Angiotensin inhibition enhances drug delivery and potentiates chemotherapy by decompressing tumour blood vessels. Nat. Commun. 4, 2516 (2013).

15. Murphy, J. E. et al. Total Neoadjuvant Therapy With FOLFIRINOX in Combination With Losartan Followed by Chemoradiotherapy for Locally Advanced Pancreatic Cancer: A Phase 2 Clinical Trial. JAMA Oncol. 5, 1020–1027 (2019).

16. Burnier, M. Telmisartan: a different angiotensin II receptor blocker protecting a different population? J. Int. Med. Res. 37, 1662–1679 (2009).

17. Wadsworth, B. J. et al. Angiotensin II type 1 receptor blocker telmisartan inhibits the development of transient hypoxia and improves tumour response to radiation. Cancer Lett. 493, (2020).

18. Mitsuhashi, A. & Nishioka, Y. Fibrocytes in tumor microenvironment: Identification of their fraction and novel therapeutic strategy. Cancer Sci. 116, 21–28 (2025).

19. Chitty, J. L. et al. A first-in-class pan-lysyl oxidase inhibitor impairs stromal remodeling and enhances gemcitabine response and survival in pancreatic cancer. Nat. Cancer 4, 1326–1344 (2023).

20. Yao, Y., et al. Pan-Lysyl Oxidase Inhibitor PXS-5505 Ameliorates Multiple-Organ Fibrosis by Inhibiting Collagen Crosslinks in Rodent Models of Systemic Sclerosis. Int. J. Mol. Sci. 23, 5533 (2022).

21. Vachhani, P. et al. PXS5505-MF-101: A Phase 1/2a Study to Evaluate Safety, Pharmacokinetics and Pharmacodynamics of Pxs-5505 in Patients with Primary, Post-Polycythemia Vera or Post-Essential Thrombocythemia Myelofibrosis. Blood 142, 625 (2023).

22. Xiao, Z. et al. Desmoplastic stroma restricts T cell extravasation and mediates immune exclusion and immunosuppression in solid tumors. Nat. Commun. 14, 5110 (2023).

23. Wang, L.-C. S. et al. Targeting Fibroblast Activation Protein in Tumor Stroma with Chimeric Antigen Receptor T Cells Can Inhibit Tumor Growth and Augment Host Immunity Without Severe Toxicity. Cancer Immunol. Res. 2, 154–166 (2014).

24. Jiang, G.-M. et al. The application of the fibroblast activation protein α-targeted immunotherapy strategy. Oncotarget 7, 33472–33482 (2016).

25. Eberlein, C. et al. A human monoclonal antibody 264RAD targeting αvβ6 integrin reduces tumour growth and metastasis, and modulates key biomarkers in vivo. Oncogene 32, 4406–4416 (2013).

26. Morris, J. C. et al. Phase I Study of GC1008 (Fresolimumab): A Human Anti-Transforming Growth Factor-Beta (TGFβ) Monoclonal Antibody in Patients with Advanced Malignant Melanoma or Renal Cell Carcinoma. PLOS ONE 9, e90353 (2014).

27. Tolcher, A. W. et al. A phase 1 study of anti-TGFβ receptor type-II monoclonal antibody LY3022859 in patients with advanced solid tumors. Cancer Chemother. Pharmacol. 79, 673–680 (2017).

28. Moore, K. M. et al. Therapeutic Targeting of Integrin αvβ6 in Breast Cancer. JNCI J. Natl. Cancer Inst. 106, dju169 (2014).

29. Anderton, M. J. et al. Induction of Heart Valve Lesions by Small-Molecule ALK5 Inhibitors. Toxicol. Pathol. 39, 916–924 (2011).

30. Faivre, S. J. et al. A phase 2 study of galunisertib, a novel transforming growth factor-beta (TGF-β) receptor I kinase inhibitor, in patients with advanced hepatocellular carcinoma (HCC) and low serum alpha fetoprotein (AFP). J. Clin. Oncol. 34, 4070–4070 (2016).

31. Melisi, D. et al. A phase II, double-blind study of galunisertib+gemcitabine (GG) vs gemcitabine+placebo (GP) in patients (pts) with unresectable pancreatic cancer (PC). J. Clin. Oncol. 34, 4019–4019 (2016).

32. Kovacs, R. J. et al. Cardiac Safety of TGF-β Receptor I Kinase Inhibitor LY2157299 Monohydrate in Cancer Patients in a First-in-Human Dose Study. Cardiovasc. Toxicol. 15, 309–323 (2015).

33. Rodon, J. et al. First-in-Human Dose Study of the Novel Transforming Growth Factor-β Receptor I Kinase Inhibitor LY2157299 Monohydrate in Patients with Advanced Cancer and Glioma. Clin. Cancer Res. 21, 553–560 (2015).

34. Sun, H., Li, T., Zhuang, R., Cai, W. & Zheng, Y. Do renin–angiotensin system inhibitors influence the recurrence, metastasis, and survival in cancer patients?: Evidence from a meta-analysis including 55 studies. Medicine (Baltimore) 96, (2017).

35. Engineer, D. R., Burney, B. O., Hayes, T. G. & Garcia, J. M. Exposure to ACEI/ARB and β-Blockers Is Associated with Improved Survival and Decreased Tumor Progression and Hospitalizations in Patients with Advanced Colon Cancer 1. Transl. Oncol. 6, 539–545 (2013).

36. Morris, Z. S. et al. Increased tumor response to neoadjuvant therapy among rectal cancer patients taking angiotensin-converting enzyme inhibitors or angiotensin receptor blockers. Cancer 122, 2487–2495 (2016).

37. Nuzzo, P. V. et al. Impact of renin-angiotensin system inhibitors on outcomes in patients with metastatic renal cell carcinoma treated with immune-checkpoint inhibitors. Clin. Genitourin. Cancer 20, 301–306 (2022).

38. Karagiannis, T., Keith, S. W., Rabinowitz, C., Louis, D. & Maio, V. Investigating Survival Associated with Angiotensin Blockade Agents in Patients with Pancreatic Cancer. Value Health 17, A71–A72 (2014).

39. Nakai, Y. et al. The inhibition of renin-angiotensin system in advanced pancreatic cancer: an exploratory analysis in 349 patients. J. Cancer Res. Clin. Oncol. 141, 933–939 (2015).

40. Wei, J. et al. Retrospective clinical study of renin-angiotensin system blockers in lung cancer patients with hypertension. PeerJ 2019, (2019).

41. Lee, C.-M. et al. Antihypertensive drugs and survival outcomes in oropharyngeal squamous cell carcinoma patients. JNCI J. Natl. Cancer Inst. djaf056 (2025) doi:10.1093/jnci/djaf056.

42. Kakuta, H., Sudoh, K., Sasamata, M. & Yamagishi, S. Telmisartan has the strongest binding affinity to angiotensin II type 1 receptor: comparison with other angiotensin II type 1 receptor blockers. Int. J. Clin. Pharmacol. Res. 25, 41–46 (2005).

43. Michel, M. C., Foster, C., Brunner, H. R. & Liu, L. A systematic comparison of the properties of clinically used angiotensin II type 1 receptor antagonists. Pharmacol. Rev. 65, 809–848 (2013).

44. Murray, C. et al. Telmisartan augments DNA damage and tumor immunogenicity to improve cancer treatments. J. Immunol. 212, 1382_5499 (2024).

45. Telmisartan Alone or Combined with Selected Standard of Care Therapy for the Treatment of Prostate Cancer - NCI. https://www.cancer.gov/about-cancer/treatment/clinical-trials/search/v?id=NCI-2024-07041 (2016).

46. UCSC Genome Browser Home. http://genome.ucsc.edu/.

47. Posit Team. RStudio: Integrated Development Environment for R. Posit Software, PBC (2023).

48. R Core Team. R: A Language and Environment for Statistical Computing. R Foundation for Statistical Computing (2025).

49. Hao, Y. et al. Dictionary learning for integrative, multimodal and scalable single-cell analysis. Nat. Biotechnol. 42, 293–304 (2024).

50. Satija, R., Hoffman, P. & Butler, A. SeuratData: Install and Manage Seurat Datasets. satijalab (2023).

51. McGinnis, C. S., Murrow, L. M. & Gartner, Z. J. DoubletFinder: Doublet Detection in Single-Cell RNA Sequencing Data Using Artificial Nearest Neighbors. Cell Syst. 8, 329–337.e4 (2019).

52. Barrett, T., et al. data.table: Extension of ‘data.frame’. 1.17.4 10.32614/CRAN.package.data.table (2025).

53. Wickham, H., François, R., Henry, L., Müller, K. & Vaughan, D. dplyr: A Grammar of Data Manipulation. (2023).

54. Bache, S. M. & Wickham, H. magrittr: A Forward-Pipe Operator for R. 2.0.3 10.32614/CRAN.package.magrittr (2022).

55. Ching, T. qs: Quick Serialization of R Objects. 0.27.3 10.32614/CRAN.package.qs (2025).

56. Bengtsson, H. R. utils: Various Programming Utilities. (2025).

57. Wickham, H. et al. Welcome to the Tidyverse. J. Open Source Softw. 4, 1686 (2019).

58. Wickham, H. Ggplot2: Elegant Graphics for Data Analysis. (Springer-Verlag New York, New York, 2016).

59. Wilke, C. O. cowplot: Streamlined Plot Theme and Plot Annotations for ‘ggplot2’. (2024).

60. Pedersen, T. L. patchwork: The Composer of Plots. (2024).

61. Garnier, S., et al. viridis: Colorblind-Friendly Color Maps for R. 0.6.5 10.32614/CRAN.package.viridis (2024).

62. Marsh, S., Tang, M., Kozareva, V. & Graybuck, L. scCustomize: Custom Visualizations & Functions for Streamlined Analyses of Single Cell Sequencing. (2024).

63. Pollard, K. S., Dudoit, S. & van der Laan, M. J. Multiple Testing Procedures: The Multtest Package and Applications to Genomics, In: Bioinformatics and Computational Biology Solutions Using R and Bioconductor. (SpringerLink, 2005).

64. Dewey, M. metap: Meta-Analysis of Significance Values. (2025).

65. Korsunsky, I., Nathan, A., Millard, N. & Raychaudhuri, S. presto: Fast Functions for Differential Expression using Wilcox and AUC. Raychaudhuri Lab (2025).

66. Aran, D. et al. Reference-based analysis of lung single-cell sequencing reveals a transitional profibrotic macrophage. Nat. Immunol. 20, 163–172 (2019).

67. Korotkevich, G. et al. Fast gene set enrichment analysis. bioRxiv 060012 (2021) doi:10.1101/060012.

68. Dolgalev, I. msigdbr: MSigDB Gene Sets for Multiple Organisms in a Tidy Data Format. (2025).

69. Pagè, H., Carlson, M., Falcon, S. & Li, N. AnnotationDbi: Manipulation of SQLite-based annotations in Bioconductor. (2024).

70. Carlson, M. org.Mm.eg.db: Genome wide annotation for Mouse. (2024).

71. Trapnell, C. et al. The dynamics and regulators of cell fate decisions are revealed by pseudotemporal ordering of single cells. Nat. Biotechnol. 32, 381–386 (2014).

72. Qiu, X. et al. Single-cell mRNA quantification and differential analysis with Census. Nat. Methods 14, 309–315 (2017).

73. Qiu, X. et al. Reversed graph embedding resolves complex single-cell trajectories. Nat. Methods 14, 979–982 (2017).

74. Cao, J. et al. The single cell transcriptional landscape of mammalian organogenesis. Nature 566, 496–502 (2019).

75. Butler, A., Hoffman, P., Satija, R. & Stuart, T. SeuratWrappers: Community-Provided Methods and Extensions for the Seurat Object. satijalab (2024).

