## Supplementary Materials for "Telmisartan alters the transcriptomes of cancer associated fibroblasts and fibrocytes in a solid tumour model"

### Slide 1
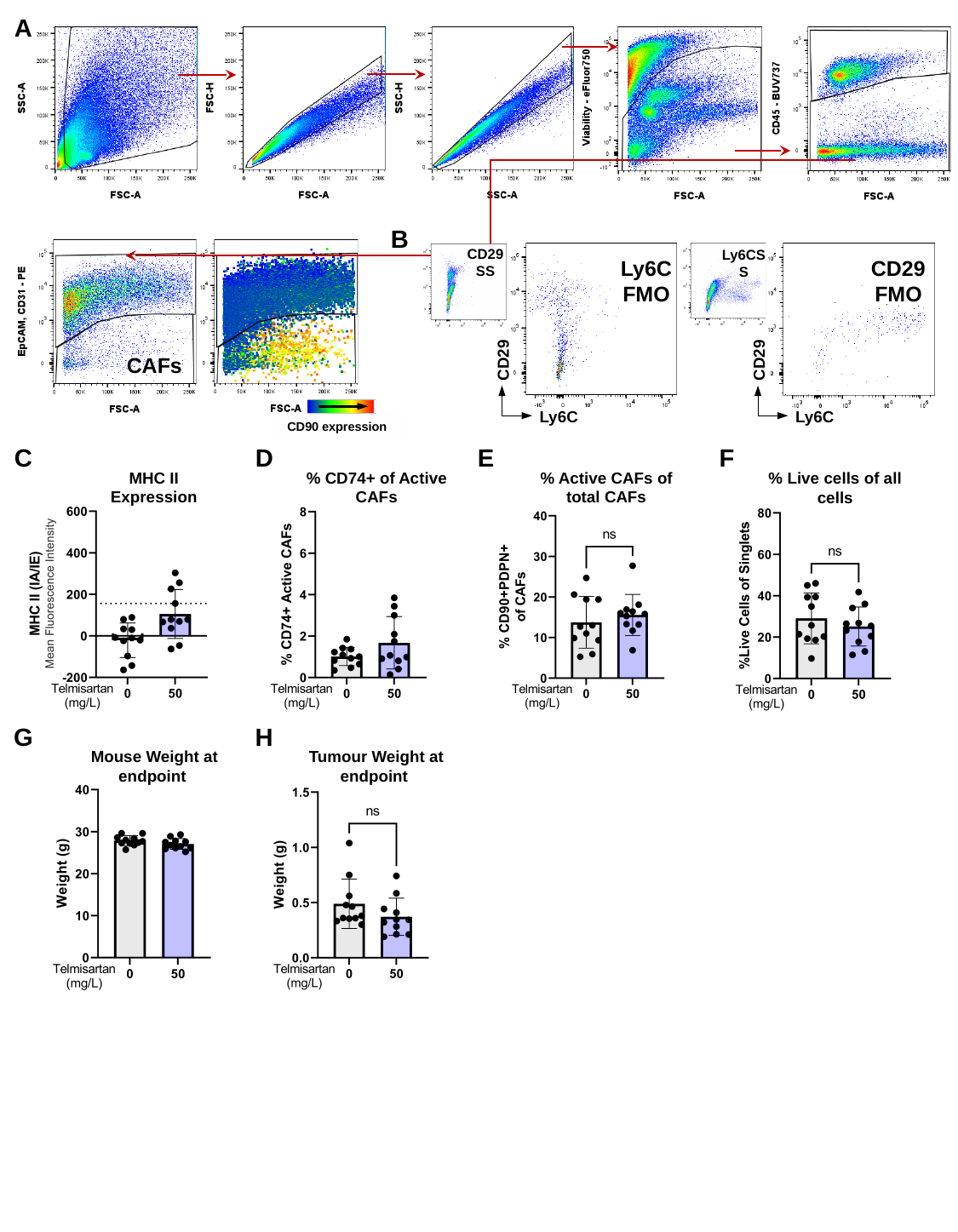

A
B
CD29 SS
Ly6CSS
Ly6C FMO
CD29
Ly6C
CD29 FMO
CD29
Ly6C
CD90 expression
CAFs
C
D
E
F
MHC II Expression
% CD74+ of Active CAFs
% Active CAFs of total CAFs
% Live cells of all cells
G
H
Mouse Weight at endpoint
Tumour Weight at endpoint

### Slide 2
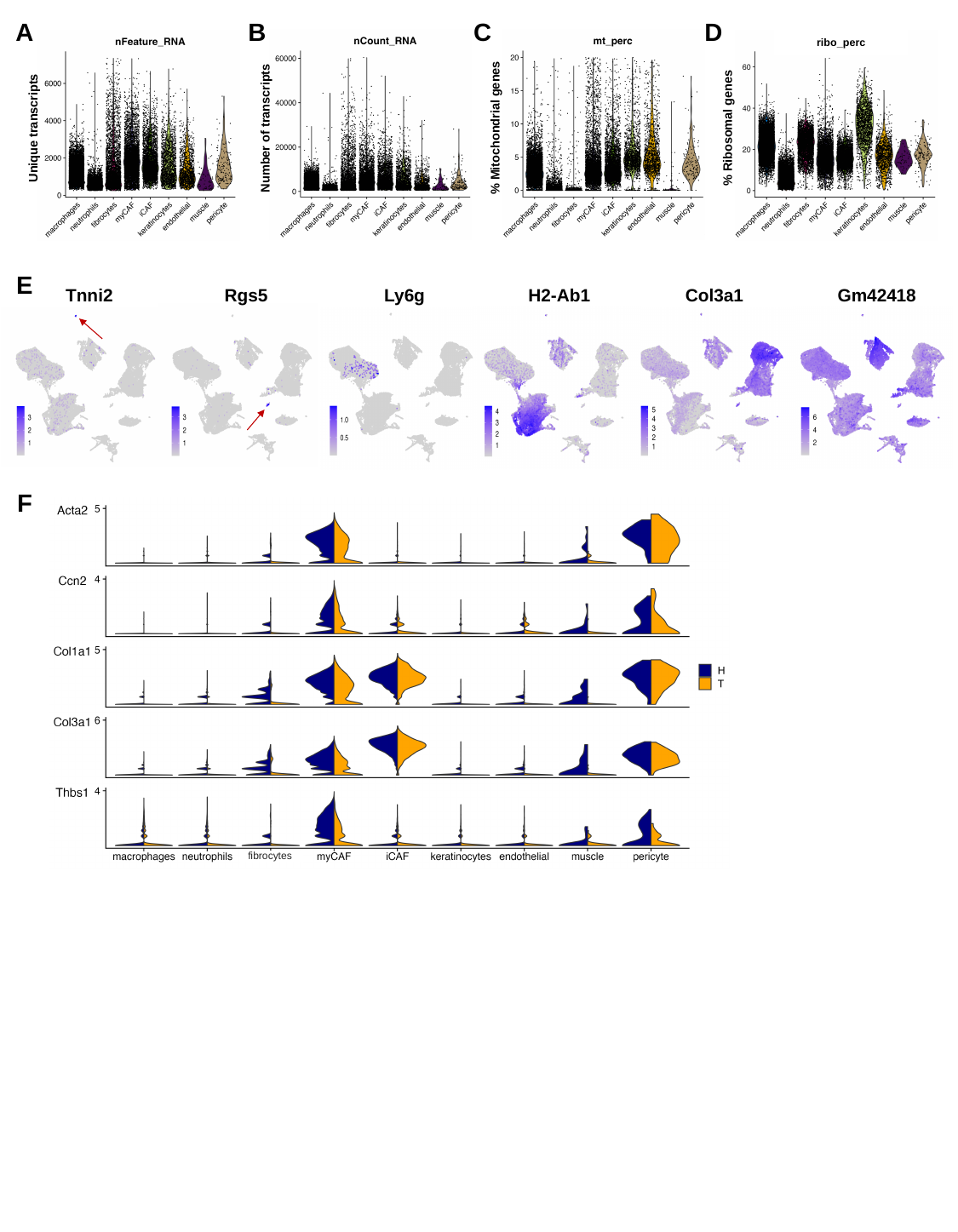

A
B
C
D
Number of transcripts
% Mitochondrial genes
ribo_perc
% Ribosomal genes
Unique transcripts
E
Tnni2
Rgs5
Ly6g
H2-Ab1
Col3a1
Gm42418
F
fibrocytes

### Slide 3
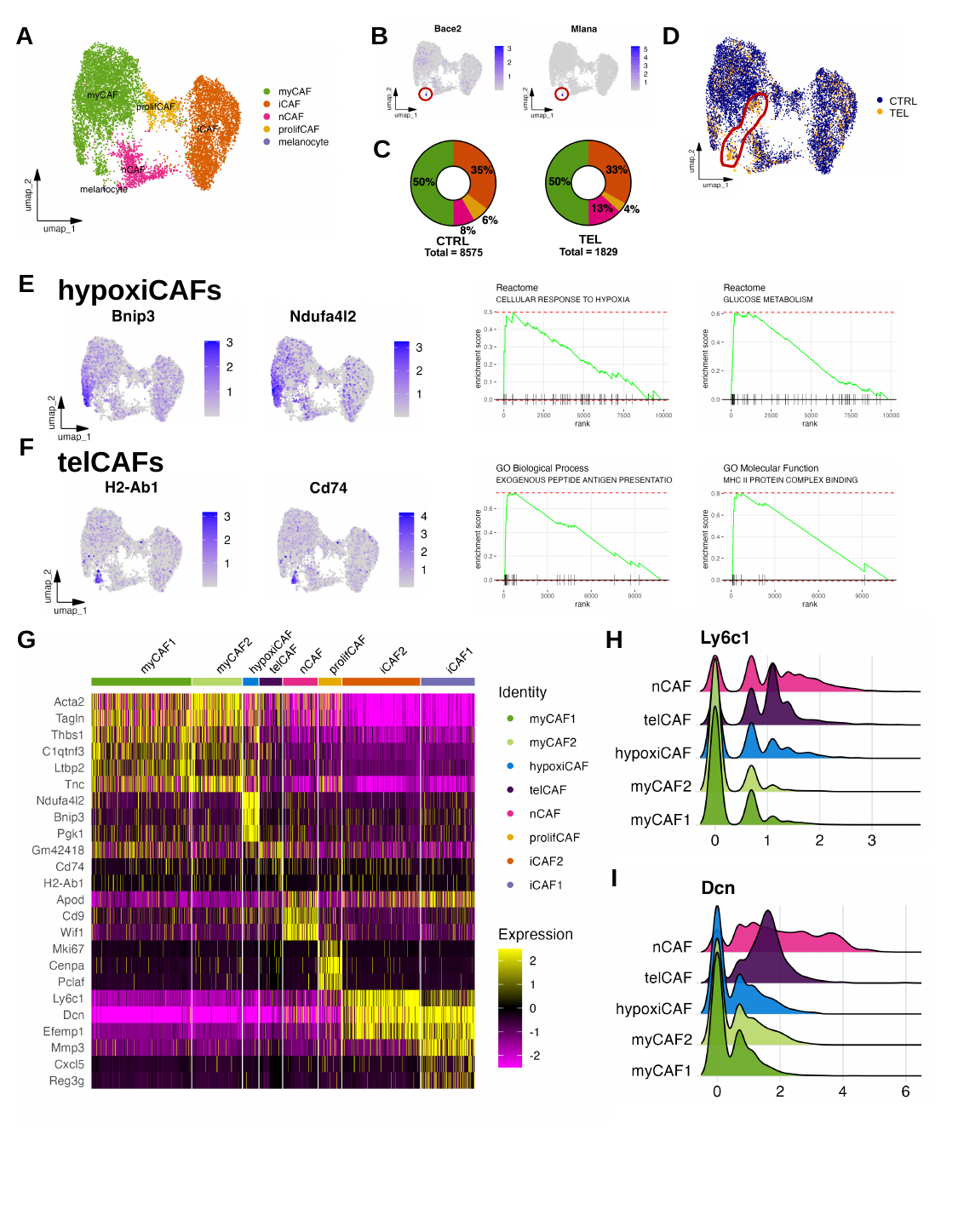

A
B
D
CTRL
TEL
C
TEL
CTRL
E
hypoxiCAFs
F
telCAFs
G
H
I
